# A literature-derived knowledge graph augments the interpretation of single cell RNA-seq datasets

**DOI:** 10.1101/2021.04.01.438124

**Authors:** Deeksha Doddahonnaiah, Patrick Lenehan, Travis Hughes, David Zemmour, Enrique Garcia-Rivera, AJ Venkatakrishnan, Ramakrisha Chilaka, Apoorv Khare, Akash Anand, Rakesh Barve, Viswanathan Thiagarajan, Venky Soundararajan

## Abstract

Technology to generate single cell RNA-sequencing (scRNA-seq) datasets and tools to annotate them have rapidly advanced in the past several years. Such tools generally rely on existing transcriptomic datasets or curated databases of cell type defining genes, while the application of scalable natural language processing (NLP) methods to enhance analysis workflows has not been adequately explored. Here we deployed an NLP framework to objectively quantify associations between a comprehensive set of over 20,000 human protein-coding genes and over 500 cell type terms across over 26 million biomedical documents. The resultant gene-cell type associations (GCAs) are significantly stronger between a curated set of matched cell type-marker pairs than the complementary set of mismatched pairs (Mann Whitney p < 6.15×10^−76^, r = 0.24; cohen’s D = 2.6). Building on this, we developed an augmented annotation algorithm that leverages GCAs to categorize cell clusters identified in scRNA-seq datasets, and we tested its ability to predict the cellular identity of 185 clusters in 13 datasets from human blood, pancreas, lung, liver, kidney, retina, and placenta. With the optimized settings, the true cellular identity matched the top prediction in 66% of tested clusters and was present among the top five predictions for 94% of clusters. Further, contextualization of differential expression analyses with these GCAs highlights poorly characterized markers of established cell types, such as CLIC6 and DNASE1L3 in retinal pigment epithelial cells and endothelial cells, respectively. Taken together, this study illustrates for the first time how the systematic application of a literature derived knowledge graph can expedite and enhance the annotation and interpretation of scRNA-seq data.

## Introduction

The development of single cell transcriptomic technologies has enabled the dissection of cellular heterogeneity within complex tissue environments [1–5]. Typically, the processing workflows for such studies involve unsupervised clustering of single cells based on their gene expression profiles followed by the assignment of a cell type annotation to each identified cluster. While cell type annotation was initially performed via manual inspection of cluster-defining genes (CDGs), there have been a number of algorithms developed recently to automate this process [6–12].

Manual cell type annotation inherently relies on an individual’s knowledge of cellular gene expression profiles. For example, an immunologist may know based on their literature expertise and firsthand experience that CD19 and CD3E are specific markers of B cells and T cells, respectively. On the other hand, the existing methods for automated annotation leverage previously generated transcriptomic datasets or curated lists of cell type-defining genes to determine which cell type a newly identified cluster most closely resembles. Interestingly, it is now common for researchers to employ both manual and automated methods for cluster annotation [13–16], as one can be used to check the veracity of the other. Importantly, there remains an unmet need for tools which augment manual annotation by transparently and objectively leveraging the associations between genes and cell types which are embedded in the literature.

Rare or novel cell types which are marked by well-studied genes have been identified through scRNA-seq, such as the CFTR-expressing pulmonary ionocyte described recently [17, 18]. Conversely, scRNA-seq should also enable researchers to identify novel markers of even well characterized cell types. However, there is currently no standardized method to assess the level of literature evidence for each individual CDG during the process of manual or automated cell type assignment. As a result, this step is often foregone in practice, in favor of proceeding to downstream workflows such as differential expression, pseudotime projection, or analysis of receptor-ligand interactions.

Here, we leverage a literature derived knowledge graph to augment the annotation and interpretation of scRNA-seq datasets. We first deploy an NLP framework to quantify pairwise associations between human protein-coding genes and cell types throughout the biomedical literature contained in PubMed. We validate that these quantified gene-cell type associations (GCAs) recapitulate canonical cell type defining genes and then harness them to perform unbiased literature-driven annotation of scRNA-seq datasets. Finally, we demonstrate that integration of these literature encoded GCAs into a differential expression workflow can highlight unappreciated gene expression patterns that warrant further experimental evaluation.

## Methods

### Generation of cell type vocabulary

To create a database of cell types, we first manually curated over 300 cell types into a directed acyclic graph structure. Specifically, each unique cell type corresponds to one node in the graph, which can be connected to one or more parent nodes (i.e. broader cell type categories) and one or more child nodes (i.e. more granular subsets of the given cell type). For example, “T cell” corresponds to one node, of which “lymphocyte” is a parent node and “CD4^+^ T cell” is a child node. Where applicable, we also manually added aliases or acronyms for each cell type. We merged this manually curated cell graph with the EBI Cell Ontology graph [19–21] by mapping identical nodes to each other and preserving all parent child relationships documented in each graph. We expanded each node to include synonymous tokens by considering synonyms from the EBI Cell Ontology and Unified Medical Language System database [22] and from our own custom alias identification service. After expansion, we reviewed our updated graph to remove erroneously introduced synonyms and performed an internal graph check to ensure that each token (i.e. cell type name or synonym thereof) occurred in one and only one node. The complete cell graph is given in **Supplemental File 1**.

For the cell type annotation algorithm described subsequently, this graph was filtered to retain the 556 nodes which were strongly associated (local score ≥ 3; see description of local scores below) with at least one human protein-coding gene; these 556 nodes are subsequently referred to as “candidate cell types.” We also defined a set of 103 “priority nodes” which intend to capture major cell types or cell type categories, to which all other candidate cell types in the filtered graph are mapped. The set of all candidate cell types, along with the priority nodes to which they map, are given in **Supplemental File 2**.

### Generation of gene vocabulary

We obtained the full set of human protein-coding genes from HGNC [23] and curated potential gene synonyms from various sources including ENTREZ, UniProt, Ensembl, and Wikipedia. For specific gene families, we also manually added family-level synonyms which are not captured by synonyms curated at the single gene level. This included genes encoding the following proteins: T cell receptor subunits, immunoglobulin subunits, class II MHC molecules, hemoglobin subunits, surfactant proteins, chymotrypsinogen subunits, CD8 subunits (CD8A, CD8B), and CD3 subunits (CD3E, CD3G, CD3D, and CD247). The complete gene vocabulary is given in **Supplemental File 3**.

For the cell type annotation algorithm described subsequently, we only considered protein-coding genes which were strongly associated with at least one cell type in the literature (local score ≥ 3; see description of local scores below). Further, we excluded mitochondrially encoded genes (gene names starting with “MT-”), genes encoding ribosomal proteins (gene names starting with “RPS”, “RPL”, “MRPS”, or “MRPL”), and MHC class I genes except for HLA-G (HLA-A, HLA-B, HLA-C, HLA-E, HLA-F). These filtering steps yielded a final set of 5,113 “eligible genes” for consideration during the cell type annotation steps (indicated in **Supplemental File 3**).

### Quantification of literature associations between genes and cell types

To quantify gene-cell type associations (GCAs) in biomedical literature, we computed local scores as described in detail previously [24]. Briefly, this metric measures how frequently two tokens *A* and *B* are found in close proximity to each other (within 50 words or fewer) in the full set of considered documents (corpus), normalized by the individual occurrences of each token in that corpus. In this case, the two tokens are a gene and a cell type, and the corpus includes all abstracts in PubMed along with all full PubMed Central (PMC) articles.

To calculate the score, we first compute the pointwise mutual information between *A* and *B* as pmi*_AB_* = log_10_([Adjacency_AB_ * N_C_]/[N_A_ * N_B_]), where Adjacency_AB_ is the number of times that Token A occurs within 50 words of Token B (or vice versa), N_A_ and N_B_ are the number of times that Tokens A and B each occur individually in the corpus, and N_C_ is the total number of occurrences of all tokens in the corpus. The local score between Tokens A and B is then calculated as LS_AB_ = ln(Adjacency_AB_ + 1) / [1 + e^−(pmiAB - 1.5)^]. A local score of 0 indicates that Tokens A and B have never occurred within 50 words of each other, and a local score of 3 indicates a co-occurrence likelihood of approximately 1 in 20. Thus, we typically consider a local score greater than or equal to 3 to represent a significant literature association.

A matrix of GCAs (i.e. pairwise local scores between all genes and all candidate cell types) is provided in **Supplemental File 4**. Notably, the maximum GCA varies substantially across the set of candidate cell types (range 3.00 - 12.54; see **Supplemental Figure 1**), which reflects the fact that different cell types have been characterized to different degrees with respect to the landscape of all human genes. To account for this, we also computed “scaled GCAs” by dividing all GCAs for a given cell type by the maximum GCA for that cell type, such that the maximum normalized GCA for every cell type is equal to 1. The matrix of scaled GCAs is provided in **Supplemental File 5**.

### Curation of canonical cell type defining genes

To test the utility of literature-derived GCAs in capturing gene expression profiles, we first curated a set of cell type defining genes, i.e. genes which were used to identify cell types in previously published manually annotated scRNA-seq datasets [25–32]. The complete set of manually curated cell type defining genes is provided in **Supplemental Table 1**. We also obtained a previously curated set of cell type defining genes from the Panglao database [33]. From this database, genes were extracted that were labeled as canonical human markers with a ubiquitousness index < 0.06, human sensitivity > 0, and mouse sensitivity > 0 (**Supplemental File 6**).

Using these curated sets of defining genes, all pairwise gene-cell type combinations and their corresponding local scores (GCAs) were then classified as “matched” or “mismatched.” A matched GCA refers to the local score between a cell type and one of its defining genes (e.g. B cells and CD19); a mismatched GCA refers to the local score between a cell type and any non-defining gene for that cell type (e.g. B cells and CD3E). That is, the mismatched pairs are obtained by simply excluding matched pairs from the set of all unique pairwise combinations of the genes and cells from the matched pairs. From the manually curated set, there were 174 matched pairs and 5,678 mismatched pairs; from the Panglao database, there were 2,291 matched pairs and 154,313 mismatched pairs.

### ROC analysis of local scores to classify matched GCAs versus mismatched GCAs

To evaluate the ability of our literature derived GCAs (local scores) to classify gene-cell type pairs as matched or mismatched, we performed a Receiver Operating Characteristic (ROC) analysis and computed the area under (AUC) the ROC curve using the “pROC” package (version 1.17.0.1) in R (version 4.0.3). Briefly, we tested over 500 thresholds (sliding intervals of 0.01 starting at 0) to determine the sensitivity and specificity of local scores in classifying gene-cell types pairs as matched or mismatched. We also repeated this ROC analysis 10,000 times with randomly shuffled assignments of “matched” and “mismatched” to empirically confirm a lack of predictive power for local scores in performing this classification with the given sets of genes and cell types when labeled at random.

### Processing of scRNA-seq studies

For each individual human scRNA-seq study listed in **Supplemental File 7**, we obtained a counts matrix and metadata file (if available) from the Gene Expression Omnibus or another public data repository. We then processed each dataset using Seurat v3.0 [34] to normalize and scale the counts, and to identify cell clusters. Normalization was performed using the NormalizeData function, with the method set to “LogNormalize” and the scale factor set to 10000, such that unique molecular index (UMI) counts were converted into values of counts per 10000 (CP10K). Scaling was performed for all genes using the ScaleData function. Linear dimensionality reduction was performed using principal component analysis (PCA), and then clusters were identified using the FindNeighbors and FindClusters functions. The top principal components explaining at least 90% of the variance were used in the FindNeighbors function, and various cluster resolutions (0.25, 0.5, 1.0, 1.5, 2.0) were tested in the FindClusters functions. Cell type annotations were obtained from associated metadata files if available; otherwise, annotation was performed manually, guided by the cell types reported in the associated publication.

### Cell type annotation algorithms

To perform automated annotation of cell types (clusters) identified from scRNA-seq datasets using our literature derived knowledge graph, we performed the following steps: (1) identify the top N cluster defining genes (CDGs), (2) compute local scores between these CDGs and all candidate cell types, (3) compute local score vector norms for each candidate cell type, and (4) rank candidate cell types for annotation plausibility based on their vector norms. Each of these steps are described in detail below, and an example workflow for annotating a single cluster is illustrated schematically in **Figures 2**-**3**. The studies for which cell type annotation was performed are listed in **Table 1** [25,27–30,35–41].

**Figure 1.**
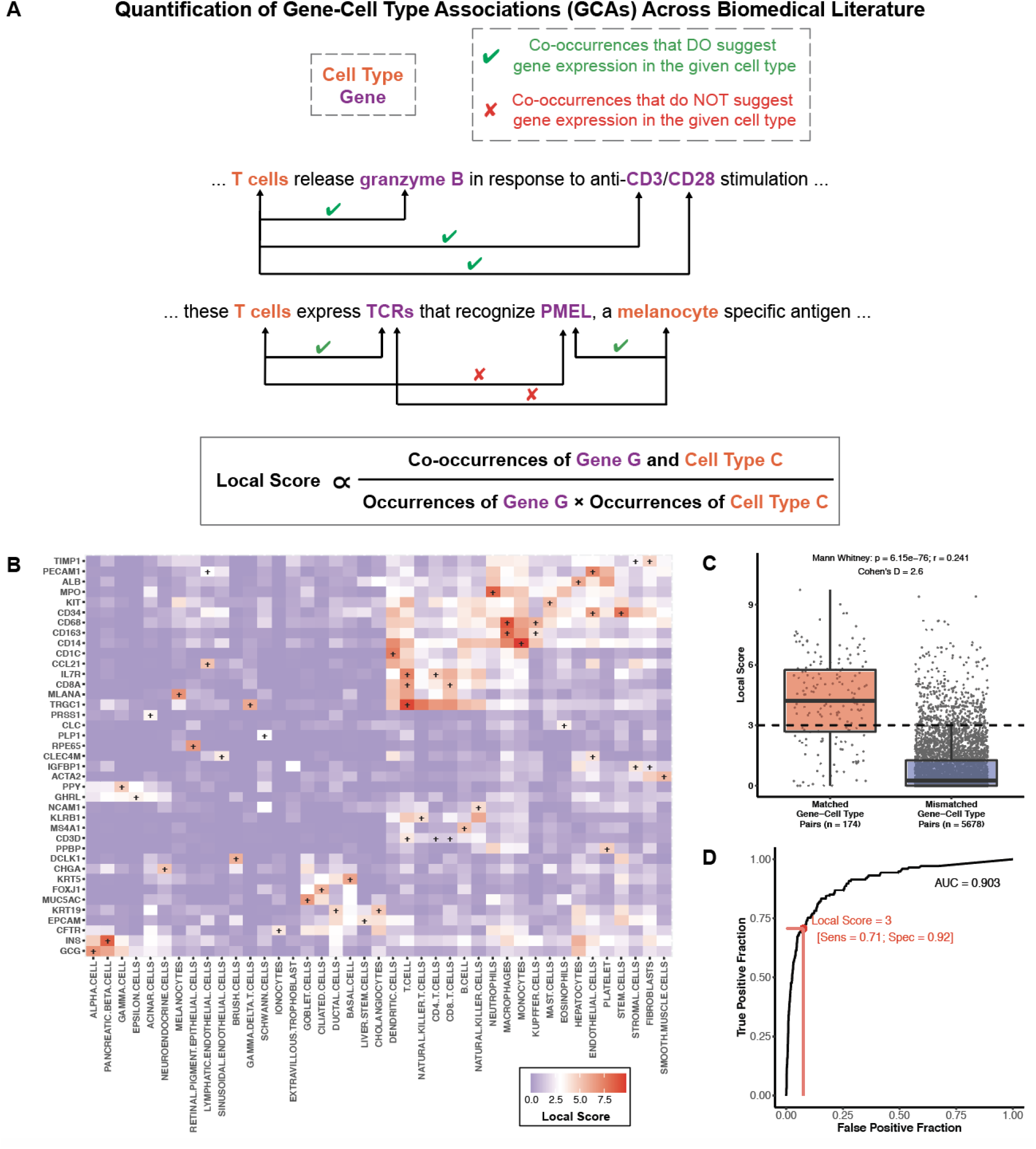
Literature derived gene-cell type associations (GCAs) capture manually curated markers of a variety of hematopoietic, epithelial, and mesenchymal cells. (A) Schematic description of the computation of local scores to quantify associations between genes and cell types across biomedical literature. The local score is a proximity metric which quantifies the likelihood of the observed co-occurrence frequency of two terms within 50 words of each other. In the context of genes and cell types, some sentences with co-occurrences state or imply that a gene is expressed in a particular cell type (denoted by green check mark), while other such sentences do not (denoted by red “X”). (B) Heatmap depicting the pairwise local scores between cell types and corresponding cell type-defining genes. These genes and cell types were extracted from a set of scRNA-seq datasets which were previously published and manually annotated. A “+” indicates a matched gene-cell type pair (i.e. genes which have been used to define the corresponding cell type in prior scRNA-seq datasets). (C) Boxplot showing the distribution of local scores (GCAs) between matched (n = 174) and mismatched (n = 5,678) gene-cell type pairs. The difference between these groups was assessed by calculating the Mann Whitney test p-value and effect size (r), along with the cohen’s D effect size. (D) Receiver operating characteristic (ROC) analysis demonstrating the ability of literature based GCAs to classify these manually curated matched versus mismatched gene-cell type pairs. The AUC was calculated as 0.904, and the sensitivity and specificity using a local score threshold of 3 are indicated in red.

**Figure 2.**
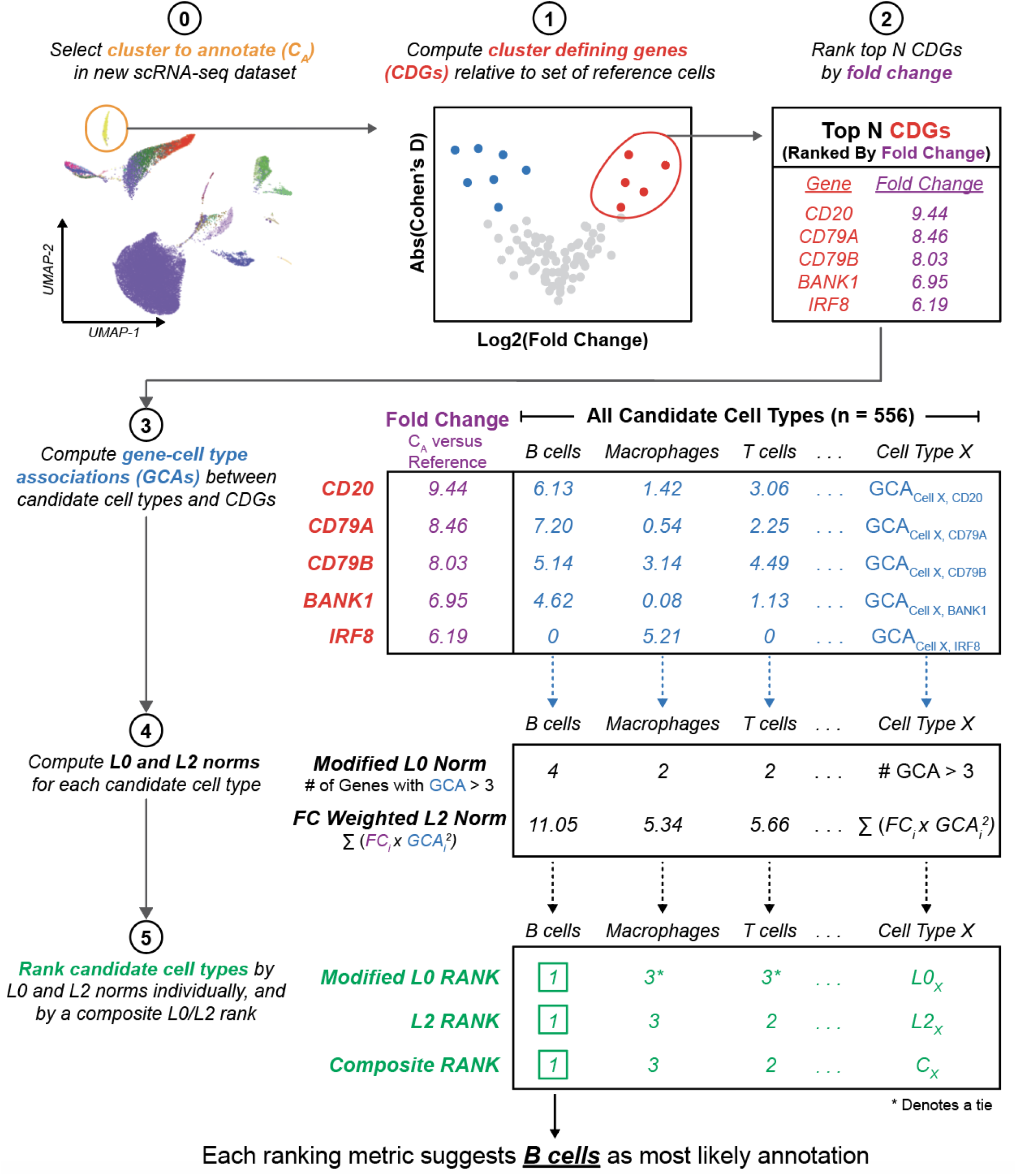
Schematic summary of the approach to leverage the literature knowledge graph to augment the annotation of cell type clusters in scRNA-seq datasets. This illustration presents the example of predicting the annotation of one cluster (B cells) from a previously published scRNA-seq dataset of the human kidney [36]. After clustering a new scRNA-seq dataset (step 1), we compute the top N cluster defining genes for a given cluster C_A_ relative to a defined set of reference cells (steps 2-3). The GCAs between each of these top N genes and all candidate cell types are computed, and the L0 and L2 norms are computed to summarize the level of literature evidence connecting these CDGs to each cell type. Candidate cell types are ranked by these norms, and the cell type showing the strongest association to the set of CDGs is selected as the most likely annotation for cluster C_A_. Note that only three candidate cell types are shown here for simplicity, but there were actually over 500 cell types considered (which mapped to 103 cell type priority nodes).

**Figure 3.**
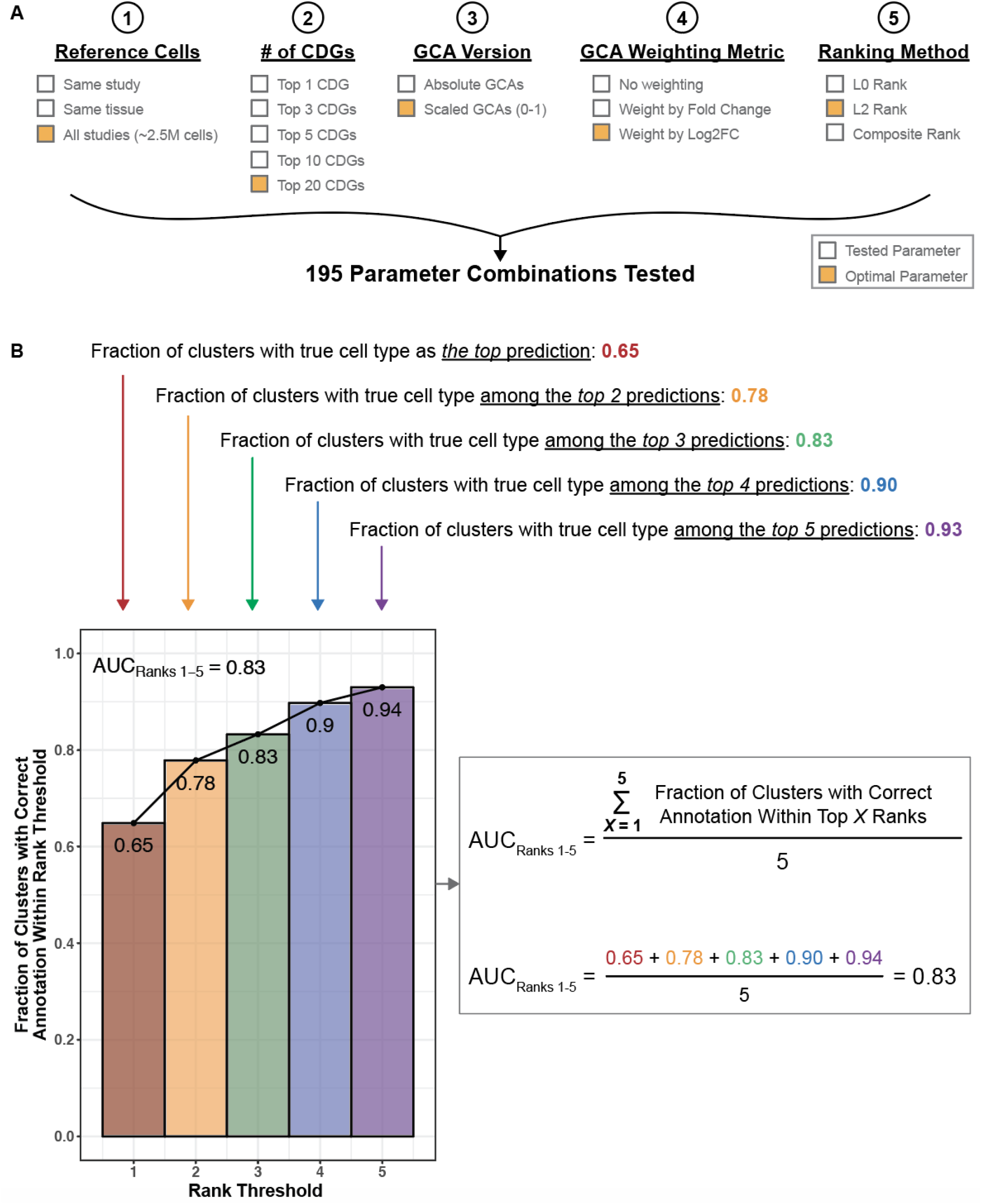
Hyperparameters tested for cluster annotation and schematic of algorithm performance evaluation. (A) Hyperparameters tested for each step of the cluster annotation algorithm, as described in Figure 2. The parameters that yielded optimal annotation performance are highlighted in orange. To compute CDGs, we compared the mean expression of all genes in the cluster of interest to their mean expression in a set of reference cells. Reference cells were taken as all other cells from the corresponding study, all other cells from the corresponding tissue, or all other cells from all processed studies. After computing fold change values for each gene, we tested the selection of 1, 3, 5, 10, and 20 genes for the downstream annotation steps. We tested the use of absolute and scaled versions of GCAs (local scores between genes and cell types). To compute L2 norms, we tested the weighting of each GCA term with the corresponding fold change and log2FC values to increase the contribution of the strongest CDGs to the cell type prediction. To rank all candidate cell types, we considered a modified L0 norm (number of genes with GCA > 3 to the given cell type), an L2 norm, and a composite metric that considers both the modified L0 and L2 norms. (B) For each parameter combination (n = 195), a cumulative distribution plot was generated to illustrate the fraction of clusters which were correctly predicted within a given rank, ranging from rank 1 to rank 5. To summarize the performance we considered the number of clusters which were annotated correctly (corresponding to the red bar at Rank Threshold = 1), and we estimated the area under this curve (denoted as AUC_Ranks 1-5_) as the average fraction of clusters for which the correct annotation was present among the top 1, 2, 3, 4, and 5 predictions.

**Table 1.**
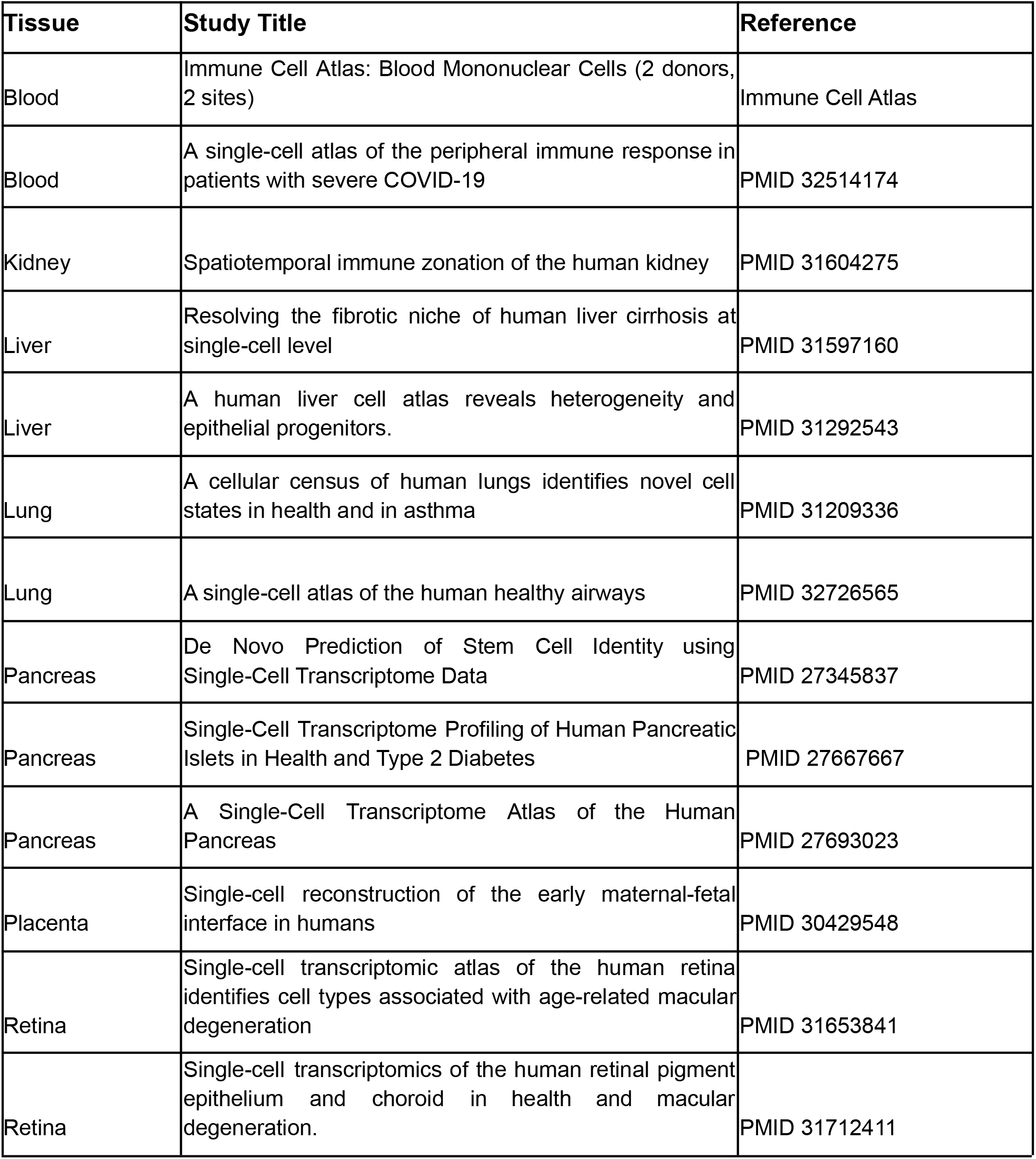
List of studies that were used to test the cluster annotation algorithm. The columns indicate (1) the tissue from which cells were derived in the study, (2) the title of the published study or dataset, and (3) the reference for the study [25,27–30,35–41]. The three pancreas datasets were integrated for one single analysis of cluster annotation.

#### 1. Identify the top N cluster defining genes (CDGs)

To compute cluster defining genes for a given cluster to annotate (*C_A_*) from study *S*, we compared the mean expression of all 5,113 eligible genes in *C_A_* to their mean expression in a reference set (*R*) of single cells. Specifically, we calculated the fold change (FC) and log2FC of mean expression for each gene, where FC = (Mean CP10K in *C_A_* + 1) / (Mean CP10K in *R* + 1) and log2FC = log2(FC).

These results were sorted in descending order and stored as two vectors: a 5,113-dimensional fold change vector *F* ([f_1_, f_2_, …, f_5113_]) and a 5,113-dimensional log2FC vector *G* ([g_1_, g_2_, …, g_5113_]).

These vectors *F* and *G* were scaled to range from 0 to 1 as follows:

- Scaled f_i_ = w_i_ = (f_i_ - F_min_) / (F_max_ - F_min_)
- Scaled g_i_ = x_i_ = (g_i_ - G_min_) / (G_max_ - G_min_)

The top N genes were then selected as CDGs, leading to the creation of two vectors for use in subsequent analyses: a N-dimensional scaled FC vector *W* ([w_1_, w_2_, …, w_N_]) and a N-dimensional scaled log2FC vector *X* ([x_1_, x_2_, …, x_N_]).

#### 2. Store absolute and scaled GCAs between these CDGs and all cell types

Absolute and scaled GCAs between the N selected CDGs for *C_A_* and all 556 candidate cell types were extracted from the GCA matrix (described above and given in **Supplemental Files 4-5**). This resulted in the generation of 556 N-dimensional vectors of absolute GCAs (one vector per candidate cell type) and 556 N-dimensional vectors of scaled GCAs (one vector per candidate cell type), represented for each candidate cell type as follows:

- Absolute GCA Vector *Y*: [y_1_, y_2_, …, y_N_]
- Scaled GCA Vector *Z*: [z_1_, z_2_, …, z_N_]

#### 3. Compute GCA vector norms for each cell type

For a given candidate cell type *C_C_*, we started with the N-dimensional vectors defined above, where each dimension corresponds to one of the N top CDGs: scaled FC (*W*), scaled log2FC (*X*), absolute GCAs (*Y*), scaled GCAs (*Z*). We then used these vectors to compute various scores quantifying the level of literature evidence connecting *C_C_* to the set of CDGs. The computed scores included variations of L0 and L2 norms as follows:

1. Modified L0 norm_Absolute GCAs_ = number of elements in *Y* greater than or equal to 3
2. L2 norm_Absolute GCAs_ = sqrt(y_1_^2^ + y_2_^2^ + … + y_N_^2^)
3. FC Weighted L2 norm_Absolute GCAs_ = sqrt(w_1_*y_1_^2^ + w_2_*y_2_^2^ + … + w_N_*y_N_^2^)
4. Log2FC Weighted L2 norm_Absolute GCAs_ = sqrt(x_1_*y_1_^2^ + x_2_*y_2_^2^ + … + x_N_*y_N_^2^)
5. L2 norm_Scaled GCAs_ = sqrt(z_1_^2^ + z_2_^2^ + … + z_N_^2^)
6. FC Weighted L2 norm_Scaled GCAs_ = sqrt(w_1_*z_1_^2^ + w_2_*z_2_^2^ + … + w_N_*z_N_^2^)
7. Log2FC Weighted L2 norm_Scaled GCAs_ = sqrt(x_1_*z_1_^2^ + x_2_*z_2_^2^ + … + x_N_*z_N_^2^)

Thus, for each candidate cell type *C_C_*, we calculated seven literature based metrics. These metrics were computed for all 556 candidate cell types, yielding a matrix of 556 candidate cell types by 7 metrics.

#### 4. Rank candidate cell types for annotation plausibility based on their vector norms

This 556 by 7 matrix was then leveraged to predict the most likely cellular identity of the cluster C_A_. Specifically, we tested the utility of each individual metric and a combination of the L0 and L2 metrics (“composite ranks”) in predicting the correct cellular identity.

When using individual metrics, cell type predictions were ranked by simply sorting the corresponding score in descending order (i.e. the prediction with the highest score was assigned a rank of 1). In the case of ties (i.e. predictions with the same score), all tied predictions were assigned the maximum (worst) possible rank; for example, if 10 predictions were tied for the highest score, then all of them were assigned a rank of 10.

To derive composite ranks, we first determined the mean and minimum of the modified L0 rank and a given L2 rank (e.g. L2 norm_Absolute GCAs_) for each cell type prediction. That is, each version of the L2 norm (n = 6) was used to generate a separate composite rank. Predictions were then ranked by sorting with the following priority order: mean rank (descending), minimum rank (ascending), and modified L0 rank (ascending). Ties were again addressed by assigning the maximum (worst) rank to all tied predictions, as described above for the handling of individual metrics.

### Hyperparameter tuning of cell type annotation algorithms

There were five adjustable parameters in our cell type annotation algorithm, which were each tested for their impact on algorithm performance as follows (see **Figure 3**):

1. *Reference set (R) of single cells used to identify the top CDGs for cluster C_A_*. We tested three options for this parameter: “within study”, “within tissue”, and “pan-study.” For the “within study” reference, the mean expression of each gene in C_A_ was compared to its mean expression in all other cells from the same study. For the “within tissue” reference, the mean expression of each gene in C_A_ was compared to its mean expression in all other cells from any study which were derived from the same tissue as C_A_. For the “pan-study” reference, the mean expression of each gene in C_A_ was compared to its mean expression in all other cells from all other processed studies (approximately 2.5 million cells; see **Supplemental File 7**). We tested these options because each has its own advantages and disadvantages. While “within study” comparisons are most commonly performed by investigators when annotating scRNA-seq datasets and are less prone to technical artifacts, selection of cell types prior to sequencing (e.g. by fluorescence activated cell sorting) can lead to the dropout of important cell type defining genes from a CDG list in this analysis workflow. For example, in a scRNA-seq study of sorted CD8^+^ T cells, important cell type markers (e.g. CD3E, CD8A, CD8B) will be ubiquitously expressed and by definition cannot be identified as CDGs for each identified subcluster. On the other hand, the “pan study” and “within tissue” comparisons are more prone to technical artifacts (e.g. batch effects, differences in sequencing depth and sample viability between studies) but are better able to preserve cell type markers in examples like the one described above. It is important to note that the pan-study comparison is also more likely to be adversely impacted by tissue contaminants (e.g. extracellular RNA) than within study or within tissue comparisons. For example, highly expressed transcripts from abundant parenchymal cells (e.g. albumin in the hepatocytes) are often detected in other non-parenchymal cells from the same tissue. When performing a pan-study comparison, this contamination of highly tissue specific transcripts could lead to the incorrect identification of these genes as markers for even the non-parenchymal cell types. However, if the cluster is compared to only other cells from the same study or tissue (which presumably have similar levels of contaminant gene expression), this issue can be avoided.
2. *Number of CDGs used to compute GCA vector norms.* We tested five options for this parameter: 1, 3, 5, 10, and 20. This range of values was selected to mirror the typical manual workflows utilized by investigators annotating their own datasets. In some cases a single obvious CDG is enough to declare a cellular identity, while in other cases it is necessary to consider the combination of several genes among the top 10 to 20 CDGs.
3. *Weighting metric used in calculating GCA L2 vector norms.* We tested three options for this parameter: no weighting, FC, and log2FC. The reason for testing this parameter is that it may be reasonable to assign more value to genes that are more strongly overexpressed in cluster C_A_ when attempting to annotate it. This hyperparameter tuning is also captured in the previous section “Compute local score vector norms for each cell type.”
4. *GCA version used to calculate L2 vector norms.* We tested two options for this parameter: absolute and scaled. The scaling of GCAs was described previously, and the raw and scaled GCAs are provided in **Supplemental Files 4-5**. This hyperparameter tuning is also captured in the previous section “Compute local score vector norms for each cell type.”
5. *Metric used to rank cell type predictions.* We tested three options for this parameter: modified L0 norm rank, L2 norm rank, and composite rank. The derivation of these ranks is described in the previous section “Rank plausible cell type annotations based on their vector norms.”

In total, we tested 195 combinations of parameters. Note that this is fewer than the total number of “possible” parameter combinations (3 x 5 x 3 x 2 x 3 = 270) because the weighting metric and GCA version used (absolute vs. scaled) in calculating L2 vector norms were irrelevant for all parameter combinations in which the modified L0 norm rank was used as the metric to rank cell type predictions.

### Performance evaluation of cell type annotation algorithms

For each cluster C_A_, our algorithm outputs a table which ranks all 556 candidate cell types, from most likely annotation to least likely annotation. We used our cell graph to map the true identity of C_A_ to its priority node (“true priority node”), and each candidate cell type was similarly mapped to its priority node (“candidate priority node”). A cell type annotation was considered correct if the true priority node was the same as the candidate priority node, and we determined the success (or failure) of labeling the given cluster by identifying the minimum (best) rank at which this was the case.

To evaluate the overall performance of each parameter combination across all surveyed studies, we first determined the fraction of clusters (out of 185 total clusters) for which the correct cell type annotation was present in the top K predictions, where K ranges from 1 to 103 (the total number of cell type priority nodes, as defined previously). For real-world application by investigators, we reasoned that an annotation would be most useful if the correct cell type was given with the top prediction, and that an annotation could also be considered useful if the correct cell type was present among the top 5 predicted cell types. To account for this definition of performance, we generated a subset of the cumulative distribution plot for each parameter combination, illustrating the fraction of clusters annotated correctly within a given rank (ranging from rank 1 to rank 5; see example in **Figure 3B**). We then estimated the area under this curve (denoted as AUC_Ranks 1-5_) by calculating the average fraction of correct cluster annotations within the top 1, 2, 3, 4, and 5 ranks (see **Figure 3B**). Parameter combinations which yielded the highest fraction of correctly annotated clusters and the highest AUC_Ranks 1-5_ were taken as the optimal algorithm settings.

### Identification of poorly characterized cell type markers

To identify potential novel or poorly characterized markers of established cell types, we compared the mean expression (CP10K) of all genes in the defined cell type of interest to their mean expression in all other cells from our reference dataset (see **Supplemental File 7**). The cell types considered here included retinal pigment epithelial cells derived from two independent studies [30, 42] and endothelial cells derived from 31 studies [25–28,30,36,38–41,43–64]. Specifically, we calculated the FC value for each gene as described above in the “pan-study” method for CDG identification during the cell type annotation algorithm. We also computed the cohen’s D (*d*) as a measure of effect size which considers the variation in the two compared groups. Specifically, this was calculated as *d* = (Mean CP10K_A_ - Mean CP10K_B_) / SD_pooled_. The pooled standard deviation was calculated as SD_pooled_ = sqrt([(N_A_-1)xSD_A_^2^ + (N_B_-1)xSD_B_^2^] / [N_A_ + N_B_ - 2]), where SD_A_ and SD_B_ are the standard deviations of each individual group, and N_A_ and N_B_ are the number of single cells in each group. Group A corresponds to the cell type of interest, and Group B corresponds to all other cells contained in the reference dataset. Among genes with *d* > 0.5, we considered the top 50 genes (ranked by fold change) as cell type markers.

After identifying the cell type markers based on transcriptional data, we assessed the literature evidence relating each marker (gene) to the cell type of interest by extracting the corresponding GCAs (local scores) from **Supplemental File 4**. Genes were classified as having Strong (local score ≥ 3.0), Intermediate (local score ≥1 and <3), or Weak (local score <1) association to the cell type of interest.

### Computation of endothelial gene signatures

After identifying potential markers of endothelial cells that have not been well characterized, we evaluated their expression levels in several datasets. To verify the given annotation of one or more clusters in each dataset as endothelial cells, we computed an endothelial signature score for each individual cell, defined as the geometric mean of CP10K values for a selected set of canonical endothelial markers; that is, Endothelial Signature = (CP10K_Gene 1_ x CP10K_Gene 2_ x … CP10K_Gene N_)^1/n^. The five endothelial markers considered were CD31 (PECAM1), AQP1, VWF, PLVAP, and ESAM. For liver sinusoidal endothelial cells, which are known to display a distinct expression profile compared to other endothelial cells, we considered CLEC4G and CLEC4M as markers rather than the previously listed set.

### Statistical analysis

Statistical analyses were performed in R (version 4.0.3). To compare local scores (GCAs) between matched and mismatched gene-cell type pairs, we computed a p-value and effect size using the two-sample Mann Whitney U Test. P-values were calculated using the *wilcox.test* function from the “stats” package (version 4.0.3), and effect sizes were calculated using the *wilcoxonR* function from the “rcompanion” package (version 2.3.27). This nonparametric test was applied because the mismatched GCAs did not show a normal distribution (**Supplemental Figures 2A-B**). We also computed cohen’s D as a measure of effect size between these groups using the *cohens_d* function from the “effectsize” package (version 0.4.3) in R. Finally, ROC curves were generated (and corresponding AUC values calculated) to assess the ability of local scores to classify matched and mismatched GCAs as described above using the “pROC” package (version 1.17.0.1).

## Results

### A literature derived knowledge graph recapitulates canonical gene-cell type associations

We have previously described the application of a knowledge graph trained on over 100 million biomedical documents to contextualize the scRNA-seq expression profile of ACE2, the entry receptor for SARS-CoV-2 [24]. Here we leverage this knowledge graph to comprehensively quantify gene-cell type associations (GCAs) across scientific publications. Specifically, we quantified each GCA as the local score between a single gene *G* and a single cell type *C* [65], which captures the likelihood of the observed co-occurrence frequency of *G* and *C* across these documents (see **Methods** and **Figure 1A**). Local scores for all such pairs are provided in **Supplemental File 4**. Of note, this metric does not account for sentiment and so will capture co-occurrences regardless of whether they denote, explicitly or implicitly, the expression of a gene in a given cell type (**Figure 1A**). Despite this potential shortcoming, we hypothesized that local scores would generally be higher for matched gene-cell type pairs (pairs of genes and cell types in which the gene is a canonical marker for given the cell type) than for mismatched pairs (pairs of genes and cell types in which the gene is not a canonical marker for the given cell type).

To test this hypothesis, we curated matched and mismatched pairs by extracting the author-provided cluster defining genes (CDGs) that were used to classify cell types in seven manually annotated scRNA-seq studies [25–32]. This yielded 174 matched gene-cell type pairs (comprising 133 unique genes and 44 unique cell types) from tissues including blood, pancreas, lung, liver, placenta, and retina (**Supplemental Table 1**). We found that indeed all cell types surveyed had a strong local association with at least one of their canonical marker genes (**Figure 1B**). For example, T cells, B cells, pancreatic beta cells, hepatocytes, and trophoblasts are strongly associated with genes including CD3E, CD20, insulin, albumin, and HLA-G, respectively (**Figure 1B**). Local scores between matched pairs were indeed significantly higher than local scores between mismatched pairs (Mann Whitney p < 6.15×10^−76^, r = 0.24; cohen’s D = 2.6; **Figure 1C**). Further, among the considered set of gene-cell type pairs, local scores were strongly predictive of whether a gene should be considered a canonical marker for a given cell type (AUC = 0.903; **Figure 1D**). We confirmed that local scores showed no predictive power when the matched and mismatched assignments were randomly shuffled (**Supplemental Figure 3**).

To confirm that these observations extend beyond the cell types and tissues captured in the studies that we curated, we performed a similar analysis to compare local scores between matched and mismatched gene-cell type pairs from the Panglao database of cell type markers [33]. From this database, we extracted 2,291 matched gene-cell type pairs (comprising 94 unique cell types and 1,666 unique genes) and 154,313 mismatched gene cell pairs (**Supplemental File 6**). We found that local scores for matched pairs were indeed significantly higher than those for mismatched pairs (Mann Whitney p < 1×10^−323^, r = 0.17; cohen’s D = 2.79; **Supplemental Figure 4A**), and that local scores robustly distinguished matched from mismatched pairs (AUC = 0.88; **Supplemental Figure 4B**). We again confirmed that local scores showed no predictive power when the matched and mismatched assignments were randomly shuffled (**Supplemental Figure 4C**).

### Literature associations facilitate augmented annotation of single cell RNA-seq datasets

Having demonstrated its ability to capture canonical cell type defining genes, we hypothesized that our knowledge graph can be applied in scRNA-seq analyses to assist in automated and unbiased cluster annotation. We thus developed an algorithm to annotate scRNA-seq datasets which have been clustered using any method of choice (see **Methods** and **Figure 2**). We tested this approach for its ability to classify 185 clusters from 13 previously annotated scRNA-seq datasets from seven human tissues: pancreas, retina, blood, lung, liver, kidney, and placenta [25–30,35–41]. This included a systematic evaluation of parameter modifications on algorithm performance (see **Methods** and **Figure 3**).

After testing the 195 possible parameter combinations, we found that optimal algorithm performance was obtained with the following settings: (1) use the pan-study reference to identify CDGs, (2) consider the top 20 CDGs, (3) use scaled local scores to compute L2 norms, (4) weight local scores by log2FC to compute L2 norms, and (5) use the L2 norm alone when ranking cell type predictions (**Figures 4A-B**; **Supplemental Table 2**). With these parameters, we were able to accurately categorize 123 of 185 (66%) cell types, and the correct annotation was among the top three or five predictions for 156 (84%) and 173 (94%) of 185 cell types, respectively (AUC_Ranks1-5_ = 0.83; **Figure 4C**; **Supplemental Table 2**). Generally, using fewer than five cluster defining genes or ranking cell type predictions by the modified L0 norm were most detrimental to algorithm performance, while the reference cell types used to compute cluster defining genes, the scaling of GCAs, and the weighting of GCAs when calculating L2 norms had less impact (**Figures 4A-B** and **Figure 5**).

**Figure 4.**
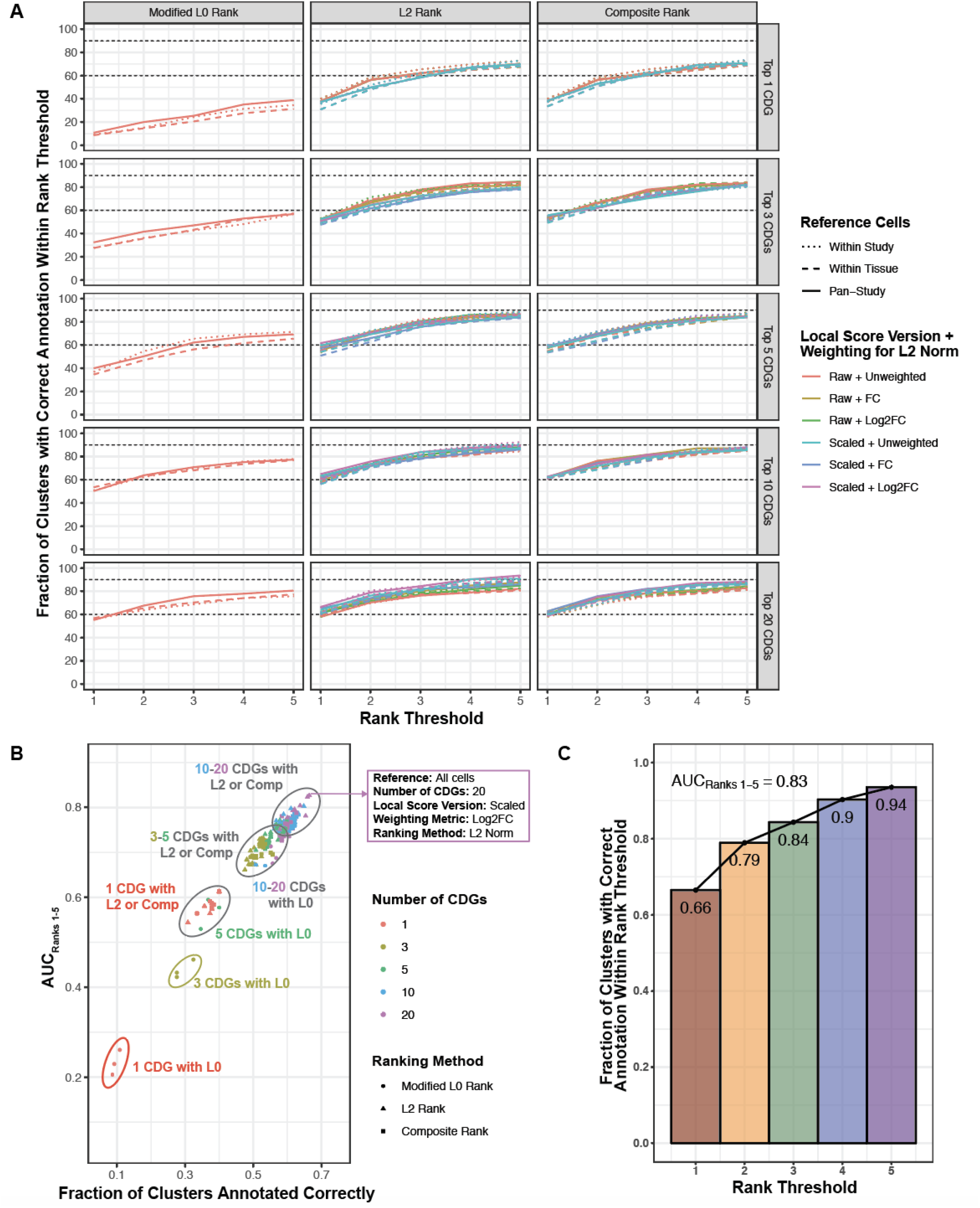
Performance evaluation of 195 parameter combinations for literature based augmented cell type annotation. (A) Each curve corresponds to one combination of parameters, where the parameters include: (1) cells used as reference to compute CDGs (encoded by line type and dot shape), (2) number of CDGs considered (encoded by the vertical facet variable), (3) the GCA version used (absolute or scaled; encoded by color), (4) the weighting method used in computed L2 norms of GCAs (also encoded by color); and (5) the metric by which candidate cell types were ranked to predict cluster identify (encoded by the horizontal facet variable). Each curve is generated from five points, specifically the percentage of clusters for which the correct annotation was present among the top 1, 2, 3, 4, or 5 predictions. (B) Performance summary of all 195 tested parametric combinations, considering the fraction of clusters which were annotated correctly (i.e. the top-ranked prediction corresponded to the true cluster label) versus the AUC_Ranks 1-5_ metric. The metrics are highly, although not perfectly, correlated. The parameter combination which showed the highest fraction of correct annotations and the highest AUC_Ranks 1-5_ was taken as the optimal approach; the optimized parameters are shown in the purple inset. (C) Summary of algorithm performance with the optimized parameters as described in (B): reference - all cells; number of CDGs - 20; local score version - scaled; weighting metric: log2FC; ranking method: L2 norm. The plot illustrates the fraction of clusters which were annotated correctly within the top 1, 2, 3, 4, or 5 ranks. The AUC_Ranks 1-5_ metric was estimated as the average of these five values.

**Figure 5.**
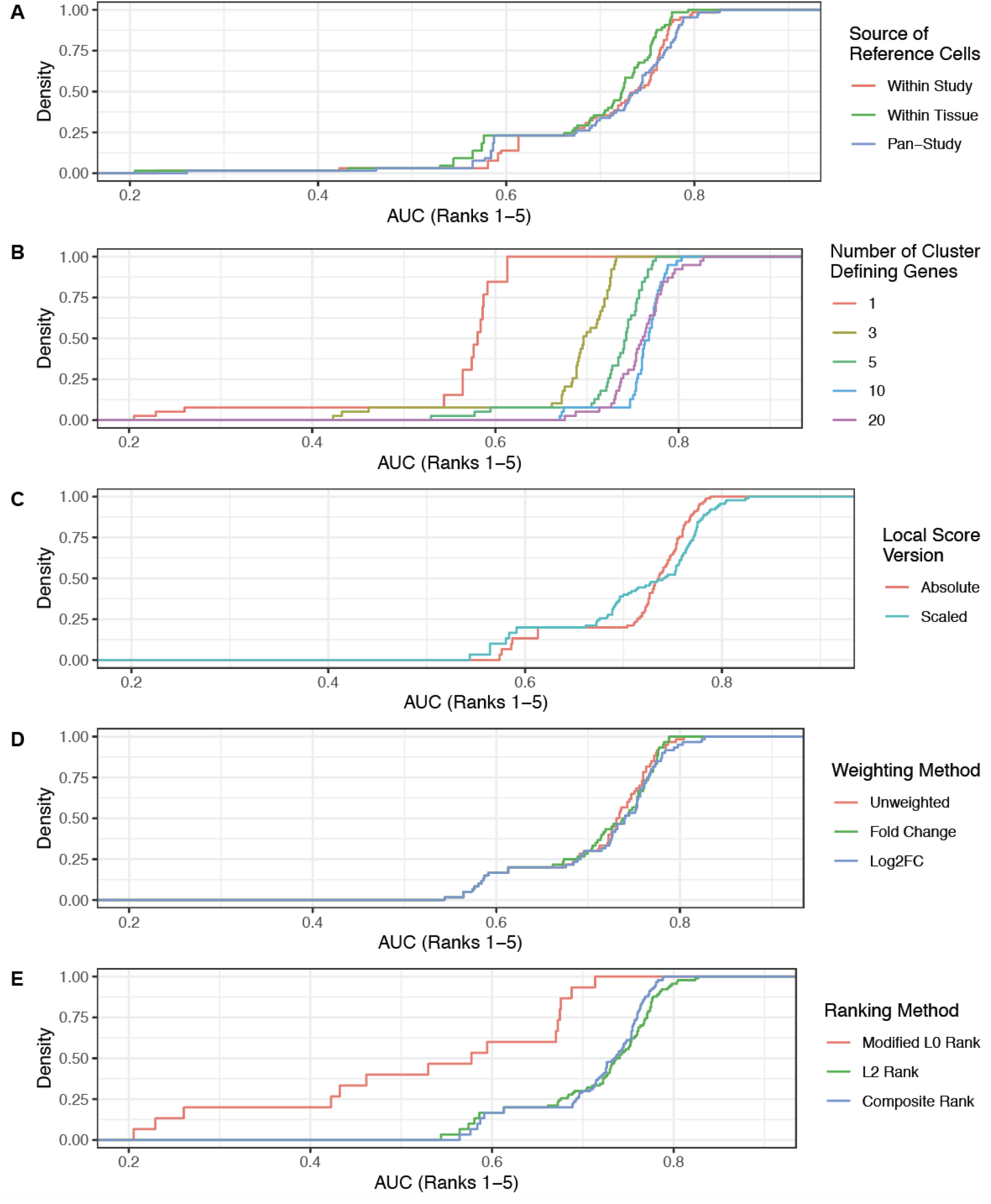
Effects of altering individual parameters on cluster annotation algorithm. Plots shown are the cumulative distribution functions of the AUC_Ranks 1-5_ metric. Each panel illustrates the effect of modifying a single parameter on the algorithm performance. (A) Modifying the source of reference cells used to compute CDGs for downstream analysis has a limited impact on performance, with the pan-study and within study options slightly outperforming the within tissue option. (B) Modifying the number of cluster defining genes considered has a strong impact on performance, with fewer genes (e.g. 1, 3, or 5) showing considerably worse performance. (C) Modifying the local score version used (absolute or scaled) has a mixed effect, with scaled GCAs contributing more of the worst-performing (left tail) and best-performing (right tailed) parameter combinations. Note that only the parameter combinations which used an L2 norm-based rank or the composite rank were included in this analysis, as only absolute GCAs were considered for the modified L0 norm-based ranking. (D) Modifying the weighting method for calculating the L2 norm has minimal impact on performance, with weighting by either fold change (FC) or log2FC slightly outperforming the unweighted method. (E) Modifying the final ranking method has a strong impact on performance, with the modified L0 rank and L2-based rank showing the worst and best performances, respectively.

The predicted annotations using our optimized algorithm parameters are shown for selected studies from retina, blood, and pancreas [25,29,30,39,40] in **Figures 6A-F**. All retinal cell types except for B cells were correctly classified, including retinal pigment epithelial cells, melanocytes, and schwann cells along with tissue resident immune and stromal cells (**Figures 6A-B**). B cells were incorrectly classified as dendritic cells, which may reflect their shared status as professional antigen presenting cells. That said, the prediction with the second highest rank was indeed B cells (**Figure 6B**).

**Figure 6.**
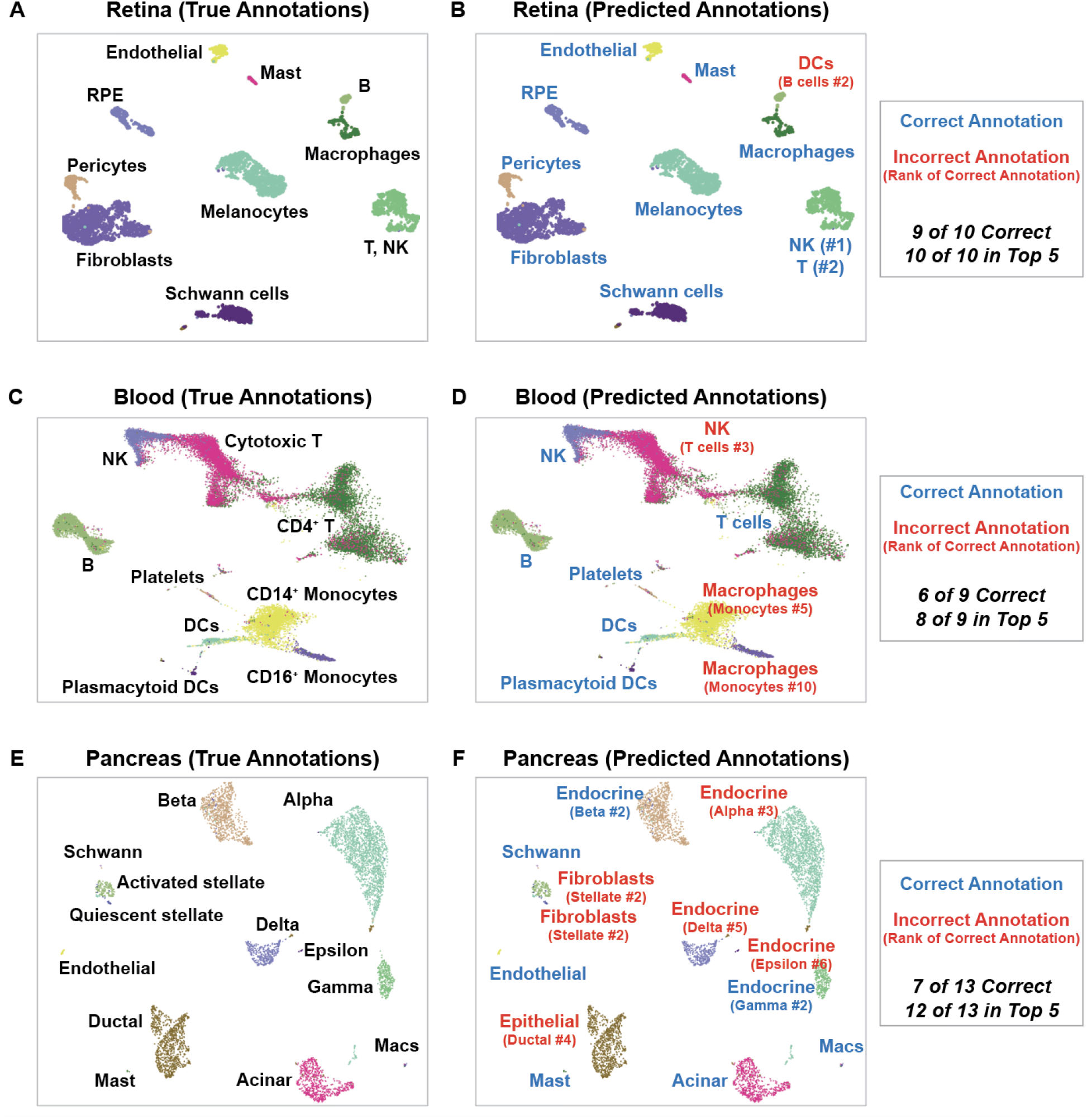
Comparison of true and predicted annotations for all cell types in three selected studies from retina, blood, and pancreas. True cluster annotations shown in the panels on the left (A, C, E) are derived from publicly deposited metadata and manual review. Predicted annotations on the right (B, D, F) are derived from our literature based cell type annotation algorithm, with the optimized parameter settings as described in the text (pan-study reference, top 20 CDGs, scaled local score, weighting by log2FC, and rank by L2 norm). Correct annotations (annotations in which the true priority node exactly matches the mapped priority node for the top prediction) are shown in green, and incorrect annotations are shown in red. For each incorrect annotation, the rank of the correct prediction (out of 103 candidate cell type priority nodes) is shown in parentheses. The datasets selected for display here were all previously published in separate studies [29,30,39,40,55].

In the blood, the labeling of monocytes proved difficult, as both CD14^+^ and CD16^+^ monocytes were misclassified as macrophages (**Figures 6C-D**). This likely reflects the close developmental and transcriptional relationships between monocytes and macrophages, as monocytes can differentiate into macrophages upon migration from circulation into tissues [66]. The misclassification of cytotoxic T cells as NK cells is also understandable, given their shared expression of cytolytic effector molecules and the fact that these cells often cluster together in scRNA-seq analyses due to their transcriptional similarity (**Figures 6C-D**).

In the pancreas, this algorithm correctly annotated acinar cells, schwann cells, endothelial cells, macrophages, and mast cells. Ductal cells were correctly classified as epithelial cells, while the specific annotation of ductal cells was ranked fourth. Alpha, beta, delta, epsilon, and gamma cells were all correctly classified as endocrine cells, with their specific subtypes ranked shortly after this broader categorization (**Figures 6E-F**). The classification of stellate cells (both quiescent and activated) as fibroblasts was deemed technically incorrect, although stellate cells are indeed known to display myofibroblast-like properties [67]. That said, stellate cells were the second ranked predictions for each of these two clusters (**Figures 6E-F**).

The predicted annotations for all other tested studies are shown in **Supplemental Figures 5-8**. It was interesting to note that certain studies were more accurately labeled with parameter settings that diverged from the overall optimized settings. For example, in one study of the retina which contained a large population of rod photoreceptors, only three of nine clusters were accurately labeled while several other cell types (e.g. amacrine cells, endothelial cells, and muller glia) were incorrectly classified as rods with the optimized settings (**Supplemental Figure 8D**). Amacrine cells were particularly problematic, with the correct label receiving a rank of 19. This suggests that many cells were contaminated with rod-specific transcripts at a high enough abundance that they dominated the CDG list when compared to all other cells in our reference set. However, this artifact was substantially mitigated by considering only the top 10 CDGs calculated using the “within-study” method and ranking predictions by the composite metric. With these settings, amacrine cells, endothelial cells, and muller glia were all annotated correctly.

### The literature knowledge graph highlights uncharacterized markers of established cell types

By contextualizing each CDG for the novelty, or lack thereof, of its association with the given cell type, our literature knowledge graph also enables researchers to rapidly identify novel markers of even well studied cell types. For example, we assessed the literature evidence for the top 50 genes overexpressed in cells of the retinal pigment epithelium (RPE) relative to all other cells in our reference dataset (**Figure 7A**). Several of these genes were established RPE markers such as RPE65 and BEST1, mutations in both of which can cause retinitis pigmentosa and other retinopathies [68, 69]. Other markers showing moderate to strong literature associations with the RPE included genes involved in vitamin A metabolism (e.g., TTR, RLBP1, RBP1, LRAT). However, we also identified several CDGs with little or no literature association to RPE cells, including genes encoding ion transporters (SLC6A13, CLIC6) and proteins that modulate Wnt signaling (FRZB, SFRP5). While these Wnt modulators have been infrequently referenced as RPE markers [70, 71], and CLIC6 has been detected by proteomics in RPE tissue and cells [72, 73], the contribution of these genes to RPE function has never been explored.

**Figure 7.**
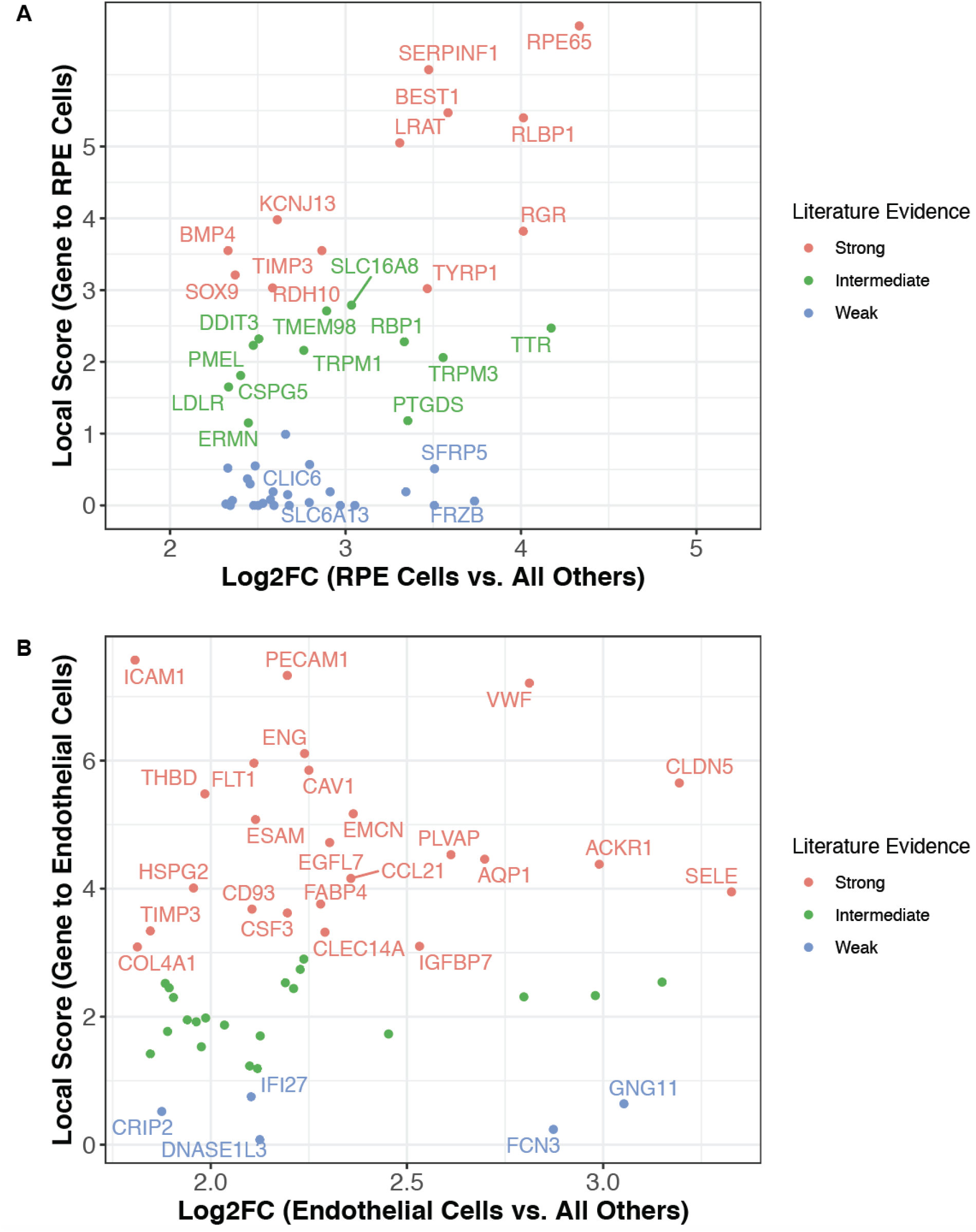
Literature derived GCAs contextualize differentially expressed genes, highlighting poorly characterized markers of established cell types. (A) Transcriptional markers of retinal pigment epithelial (RPE) cells were identified by comparing the mean expression of all genes in RPE cells from two scRNA-seq studies [30, 42] to their expression in all other cells from our reference datasets (see Supplemental File 7). The top 50 markers (ranked by fold change) were classified for their level of literature association to retinal pigment epithelial cells as follows: strong: Local Score ≥ 3; intermediate: Local Score ≥ 1 and < 3; weak: Local Score < 1. All of the genes with strong or intermediate evidence are highlighted by name, as are a subset of the genes with weak evidence that may warrant further evaluation. (B) The same process was applied as in (A), but for endothelial cells from 31 studies [25–28,30,36,38–41,43–64] rather than for RPE cells. Here, all of the genes with strong or weak evidence are highlighted by name.

Similarly, we assessed the existing literature evidence for genes overexpressed in endothelial cells (ECs). While 24 of the top 50 genes were strongly associated with ECs (e.g., PECAM1, VWF, ICAM1), several other genes were poorly characterized or previously uncharacterized endothelial markers (**Figure 7B**). For example, DNASE1L3 was identified as an EC marker whereas it is canonically reported to be expressed by macrophages and dendritic cells [74–76]. While its expression in liver sinusoidal ECs, non-sinusoidal hepatic ECs, and renal ECs of the ascending vasa recta has been recently reported [57,77–79], we not only confirmed expression in these populations (**Supplemental Figure 9**) but also identified several other tissues in which ECs were the predominant DNASE1L3-expressing cell type, including the adrenal gland, lung, and nasal cavity (**Supplemental Figure 10**). Further, the functional significance of DNASE1L3 expression in ECs has not been explored, but it may indeed be very relevant given the strong genetic associations connecting DNASE1L3 to the development of anti-dsDNA antibodies and various autoimmune phenotypes including lupus [75,76,80], systemic sclerosis [81], scleroderma [75, 80], and hypocomplementemic urticarial vasculitis syndrome [82].

## Discussion

scRNA-seq has enabled the characterization of cellular heterogeneity at unprecedented levels. Initial uptake was relatively slow due to the costs associated with these experiments, but rapid technological advances have drastically improved the accessibility of this technique [83]. As a result, the amount of scRNA-seq data which is being generated and deposited into public databases is likely to continue increasingly rapidly for years to come. It is imperative that investigators are equipped to efficiently annotate and analyze these datasets such that the insights embedded within them are not left untapped.

The cell type annotation tools developed in recent years provide excellent resources that will assist in efforts to analyze individual datasets and to synthesize the exponentially increasing quantity of publicly available data [6–12]. However, these methods do have intrinsic shortcomings, such as the requirement for the existence or generation of high quality reference transcriptomic datasets for all cell types that may be recovered in a given scRNA-seq experiment. By considering all cell types that have been described in the literature, our method circumvents this need for data curation. Further, even as these tools are increasingly applied to automate data processing pipelines, it remains preferable to perform some degree of manual review to ensure the validity of the resultant annotations. Specifically, the quality of annotations should minimally be assessed by determining whether a set of CDGs is biologically consistent, per canonical knowledge, with the given cell type label. Our method is specialized to perform this exact task.

Standard manual annotation is flawed due its inherent subjectivity and reliance on the existing knowledge of a single investigator. These flaws can be mitigated to some degree by “assisted” manual annotation that leverages search engines (e.g., Google, PubMed) to identify cell types which have been reported to express the genes under consideration. However, this process only leverages a sliver of the information contained within the biomedical literature and would be prohibitively time consuming if scaled to the analysis of large numbers of datasets. On the other hand, the deployment of NLP algorithms to mimic manual annotation provides a superior alternative, enabling “augmented annotation” workflows that objectively harness the entire knowledge graph of GCAs contained in the literature to suggest which cell types are most strongly associated with a given set of CDGs. By integrating this approach with already existing annotation workflows, one can rapidly assess the veracity of predicted cell type annotations and highlight those which warrant further review by an expert.

The utility of this literature knowledge graph also extends beyond the annotation of cell types, unlocking a new analytic workflow in the interpretation of scRNA-seq data. By its very nature, scRNA-seq provides investigators with data at the level of gene-cell type pairs, i.e. *Gene X is expressed in Cell Type Y at Level Z*. However, to date there has been no systematic and objective method to comprehensively contextualize this data with respect to the world’s knowledge of those same gene-cell type pairs at that point in time. To address this unmet need, we demonstrated that our database of literature derived GCAs can be used to assess the degree of literature evidence connecting any gene to any cell type in which its expression has been observed by scRNA-seq. This workflow will provide investigators with the opportunity to rapidly identify gene expression patterns which were previously unknown or poorly characterized, and thereby prioritize candidates for directed follow-up functional studies.

For example, we found CLIC6 and DNASE1L3 as uncharacterized markers of RPE cells and endothelial cells, respectively. Although CLIC6 was identified almost two decades ago as a member of the chloride intracellular channel (CLIC) gene family, its function remains essentially unknown [84, 85]. Biochemical studies have demonstrated that CLIC6 interacts with dopamine D2 receptors in the brain, but it is still unclear whether CLIC6 modulates dopamine receptor mediated signaling or even serves as a functional chloride channel [85, 86]. It is intriguing that CLIC4, another member of the CLIC family, functions in RPE cells to promote their epithelial morphology, maintain their attachment to the photoreceptor layer of the retina, and regulate extracellular matrix degradation at focal adhesions [87, 88]. CLIC6, on the other hand, has never been functionally characterized in these cells despite having been detected in them by proteomics and immunohistochemistry [72,73,89]. Given recent evidence for structural conservation between CLIC6 and other CLIC family proteins [90], we suggest that directed studies to analyze CLIC6 function in RPE cells are warranted.

DNASE1L3 is an extracellular DNase which is historically reported to be released specifically by macrophages and dendritic cells [74,75,80]. Our analysis challenges this canon, insteading highlighting this gene as an uncharacterized marker of endothelial cells (in addition to myeloid cells) in tissues including liver, kidney, lung, nasal cavity, thyroid, and adrenal cortex. Interestingly, DNASE1L3 is genetically associated with multiple autoimmune phenotypes including systemic lupus erythematosus (SLE) [75, 80], systemic sclerosis [81], and scleroderma [75, 80]. Loss of function mutations in DNASE1L3 are responsible for a familial form of SLE which is characterized by the presence of anti-dsDNA antibodies and lupus nephritis [76], and mechanistic studies have directly implicated DNASE1L3 deficiency as a cause of anti-dsDNA antibody development [75]. Further, DNASE1L3 mutations have been reported to cause hypocomplementemic urticarial vasculitis syndrome, an inflammatory disease of the vascular system which often progresses to SLE [82]. Given these genetic associations, we hypothesize that the production of DNASE1L3 by ECs may protect against the development of autoantibodies and outright autoimmune disease. While CD11c^+^ cells are responsible for about 80% of serum DNASE1L3 activity in mice [75], it is plausible that ECs contribute significantly to the remaining 20% or that EC-derived DNASE1L3 acts in a more localized fashion. Indeed, the localized activity of other EC products, such as tissue-type plasminogen activator (t-PA) and von Willebrand factor (VWF), are known to regulate clot formation specifically at sites of damaged endothelium. Perhaps the activity of DNASE1L3 in the close vicinity of renal, hepatic, and pulmonary endothelial cells protects against the development of nephritis and other forms of vasculitis at these sites.

There are several limitations of this study. First, the GCAs which are used to perform cluster annotation simply measure literature proximity of a gene and a cell type without accounting for the sentiment surrounding this co-occurrence. The algorithm would be improved by training NLP models which can distinguish between co-occurrences that denote a gene expression relationship versus those that are ambiguous, spurious, or explicitly deny a gene expression relationship. Second, annotation using literature derived associations inherently has limited utility in the recognition of novel cell types and cell types which are infrequently discussed in literature. Third, the granularity derived from this method is lower than that derived from many other existing pipelines which use reference transcriptomes of defined cellular subsets to automate the annotation process.

With these shortcomings in mind, we highlight that this should be viewed as a tool for augmenting current annotation workflows rather than as a standalone automated pipeline to replace other methods. In the future, formal integration of our literature based method with data-driven annotation tools will reduce the time and effort required for manual review of automated annotations. Further, we emphasize that the utility of quantified literature associations in scRNA-seq analyses extends beyond cluster annotation, enabling for the first time a rapid, systematic, and unbiased novelty assessment of all observed gene expression patterns from any previously or newly generated dataset.

Taken together, we have presented a new framework for the processing and interpretation of scRNA-seq datasets. Using a literature derived knowledge graph, we comprehensively quantified the strength of associations between human genes and cell types. These associations robustly capture relationships between many cell types and their canonical gene markers, and accordingly they can be used to annotate clusters of distinct cell types identified by scRNA-seq. Finally, these associations can rapidly contextualize lists of CDGs and differentially expressed genes, enabling investigators to identify and prioritize uncharacterized cell type markers for further exploration.

## Acknowledgements

We thank Murali Aravamudan for his thoughtful review and feedback on this manuscript.

## Data Availability

The datasets used to test the methods presented were accessed from the NCBI Gene Expression Omnibus (GEO) or another public data repository (see **Table 1** and **Supplemental File 7**). These datasets, in the processed and annotated form as used in our analyses, can be accessed and downloaded by academic researchers from academia.nferx.com, and will be made accessible to non-academic researchers upon reasonable request. All newly generated data used in the cluster annotation workflow (e.g. cell graph, cell type mapping to priority nodes, raw and scaled GCA matrices) are available in **Supplemental Files 1-5**, and all code used to perform and evaluate cluster annotations will be made available on github.

## Author Contributions

*Deeksha Doddahonnaiah*: Data curation, Formal analysis, Investigation, Methodology, Validation, Visualization, Writing - original draft, Writing - review and editing. Contributed equally with Patrick Lenehan.

*Patrick Lenehan*: Conceptualization, Formal analysis, Supervision, Investigation, Methodology, Validation, Visualization, Writing - original draft, Writing - review and editing. Contributed equally with Deeksha Doddahonnaiah.

*Travis Hughes*: Conceptualization, Data curation, Formal analysis, Investigation, Methodology, Writing - review and editing.

*David Zemmour*: Conceptualization, Data curation, Formal analysis, Investigation, Methodology, Writing - review and editing.

*Enrique Garcia-Rivera*: Conceptualization, Methodology, Writing - review and editing.

*AJ Venkatakrishnan*: Writing - review and editing.

Ramakrisha Chilaka: Software.

Apoorv Khare: Software.

*Akash Anand*: Software.

*Rakesh Barve*: Resources, Supervision, Project administration.

*Viswanathan Thiagarajan*: Resources, Supervision, Project administration.

*Venky Soundararajan*: Conceptualization, Resources, Supervision, Funding acquisition, Validation, Investigation, Methodology, Project administration, Writing - review and editing.

## Supplemental Figures

**Supplemental Figure 1.**
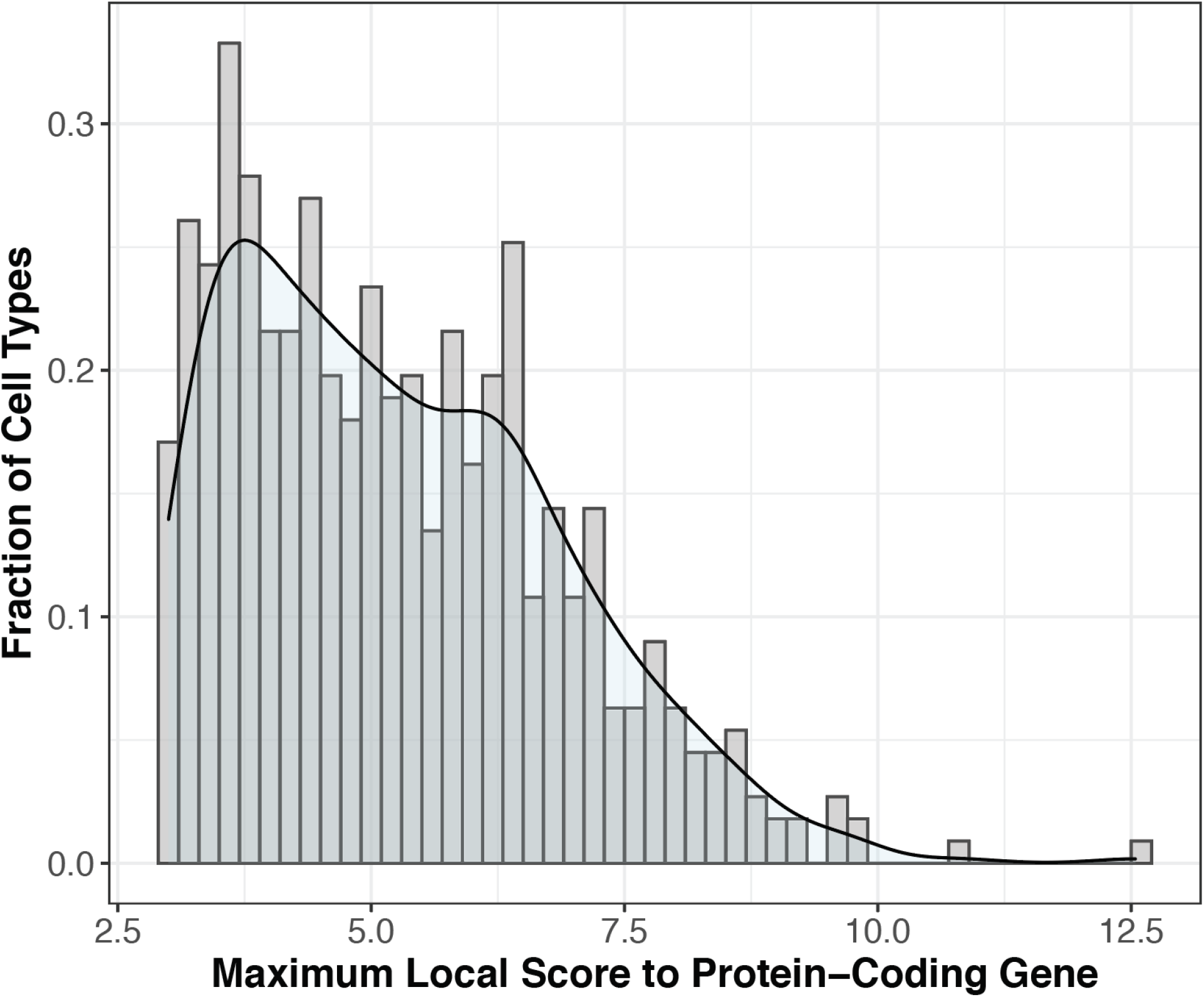
Distribution of maximum GCAs for all 556 candidate cell types. For each candidate cell type (n = 556), the maximum GCA (local score between any human protein-coding gene and the given cell type) was extracted from **Supplemental File 3**. This distribution shows that the maximum GCA varies substantially by cell type (range 3.00 - 12.54), and so we also scaled these values to range from 0 to 1 for each cell type (see **Supplemental File 4**). Both absolute and scaled GCAs were tested for their utility in cluster annotation.

**Supplemental Figure 2.**
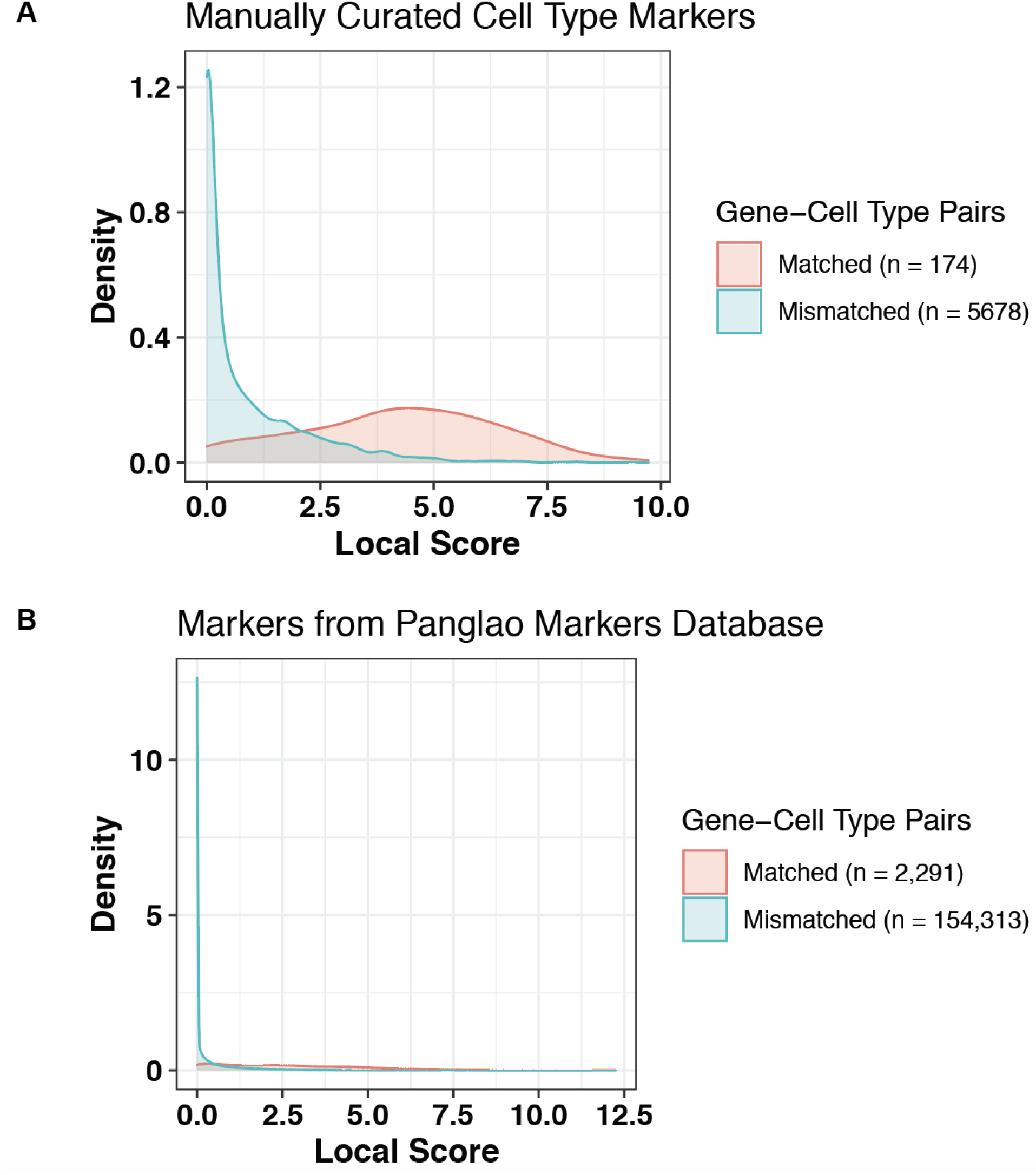
Distribution of gene-cell type local scores (GCAs) between matched and mismatched pairs. A “matched” pair corresponds to (A) a gene which was used to define a cell type in prior scRNA-seq analyses and its corresponding cell type, or (B) a gene which are documented as canonical human cell type markers in the Panglao database and its corresponding cell type [33]. In each case, after obtaining the set of matched pairs, all other possible gene-cell type combinations (i.e. all other pairwise combinations of these genes and cell types) were considered “mismatched” pairs. The distributions here show that these local scores do not follow a normal distribution, and so we used nonparametric tests to assess the difference between them.

**Supplemental Figure 3.**
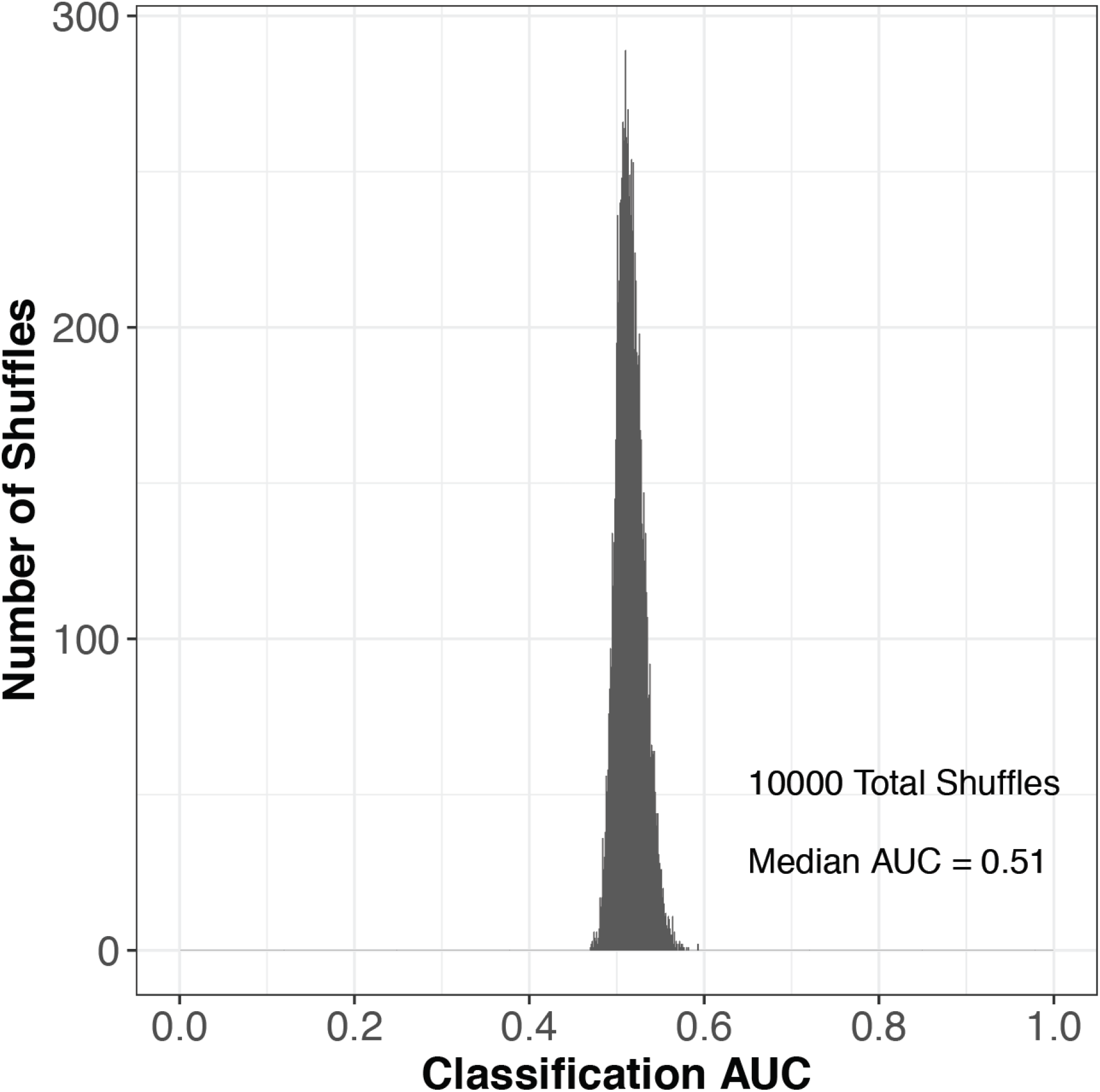
Distribution of AUC values from ROC analysis of GCA-based classification of manually curated gene-cell type pairs with randomly shuffled matched and mismatched assignments. A set of matched gene cell-type pairs was manually curated, and the complement pairs were designated as mismatched. After performing an ROC analysis to determine the classification power of GCAs in discriminating matched from mismatched gene-cell type pairs, we performed 10,000 iterations of this ROC analysis with random shuffling of the matched and mismatched labels. This histogram shows the distribution of the 10,000 AUC values obtained from these analyses.

**Supplemental Figure 4.**
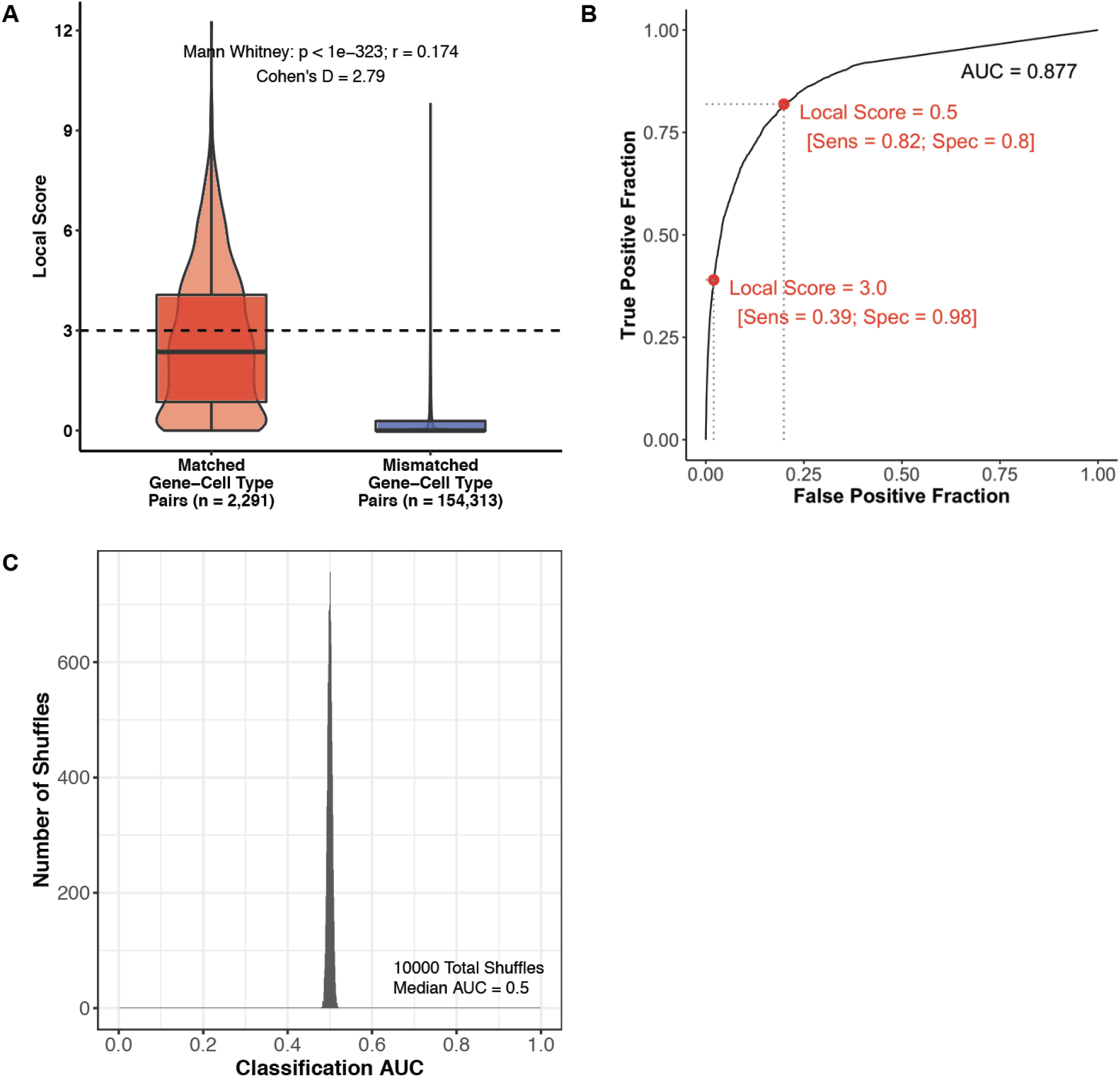
Literature based GCAs distinguish matched from mismatched gene-cell type pairs derived from the Panglao database of cell type markers. (A) Boxplot and violin plot showing the distribution of local scores (GCAs) between matched (n = 2,291) and mismatched (n = 154,313) gene-cell type pairs. The difference between these groups was assessed by calculating the Mann Whitney test p-value and effect size (r), along with the cohen’s D effect size. (B) Receiver operating characteristic (ROC) analysis demonstrating the ability of literature based GCAs to classify matched versus mismatched gene-cell type pairs. The AUC was calculated as 0.877, and the sensitivity and specificity at specific local score thresholds are indicated in red. (C) Distribution of AUC values from 10,000 repeats of the analysis in (B), where the matched and mismatched labels were randomly shuffled prior to performing the ROC analysis. As expected, GCAs do not show any classification power when the labels are randomly assigned.

**Supplemental Figure 5.**
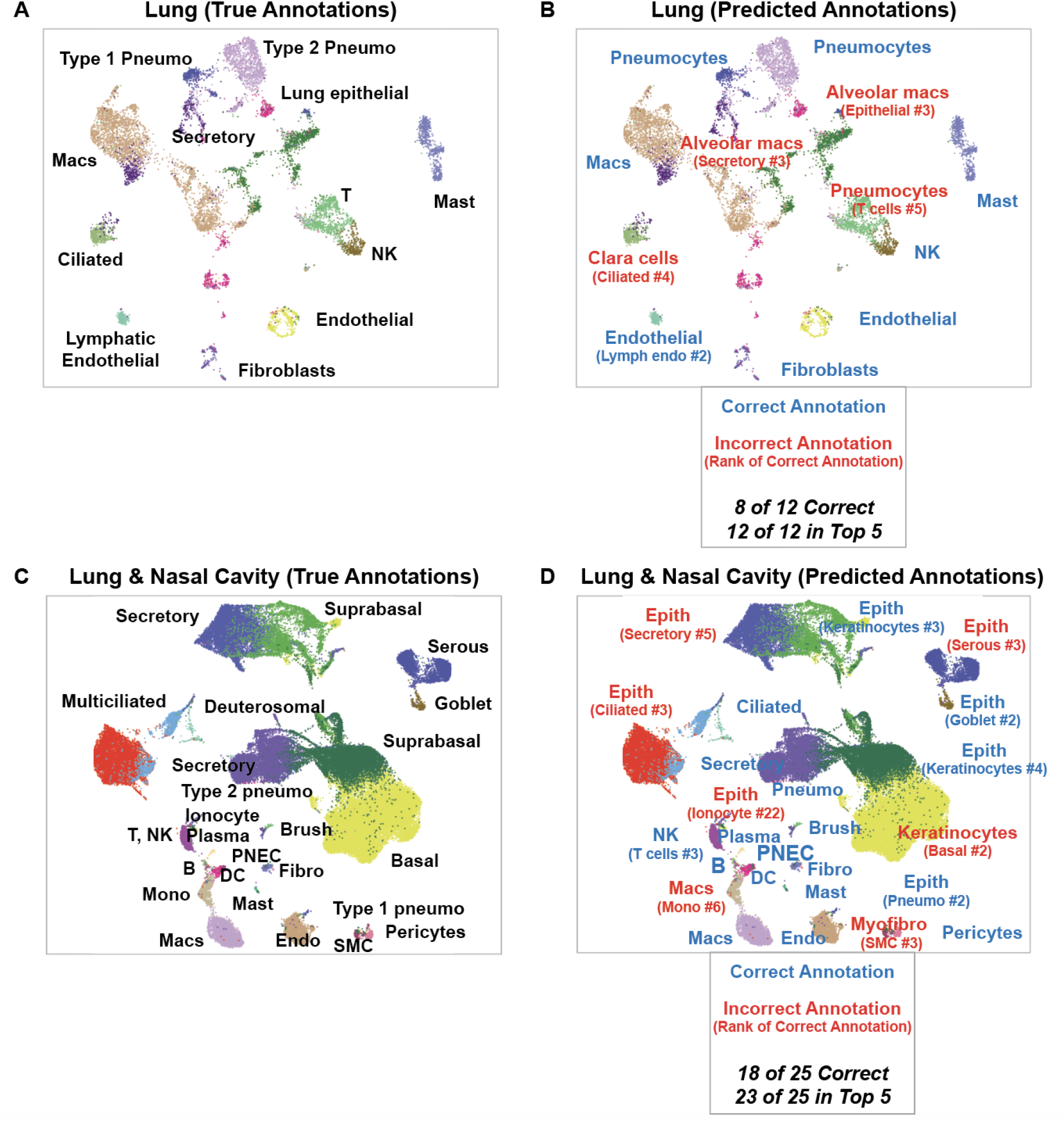
Comparison of true and predicted annotations for all cell types from two scRNA-seq studies of the respiratory tract (lung and nasal cavity). True cluster annotations shown in the panels on the left (A, C) are derived from publicly deposited metadata and manual review. Predicted annotations on the right (B, D) are derived from our literature based cell type annotation algorithm, with the optimized parameter settings as described in the text (pan-study reference, top 20 CDGs, scaled local score, weighting by log2FC, and rank by L2 norm). Correct annotations (annotations in which the true priority node exactly matches the mapped priority node for the top prediction) are shown in green, and incorrect annotations are shown in red. For each incorrect annotation, the rank of the correct prediction (out of 103 candidate cell type priority nodes) is shown in parentheses. The datasets selected for display here were previously published in separate studies [26, 38].

**Supplemental Figure 6.**
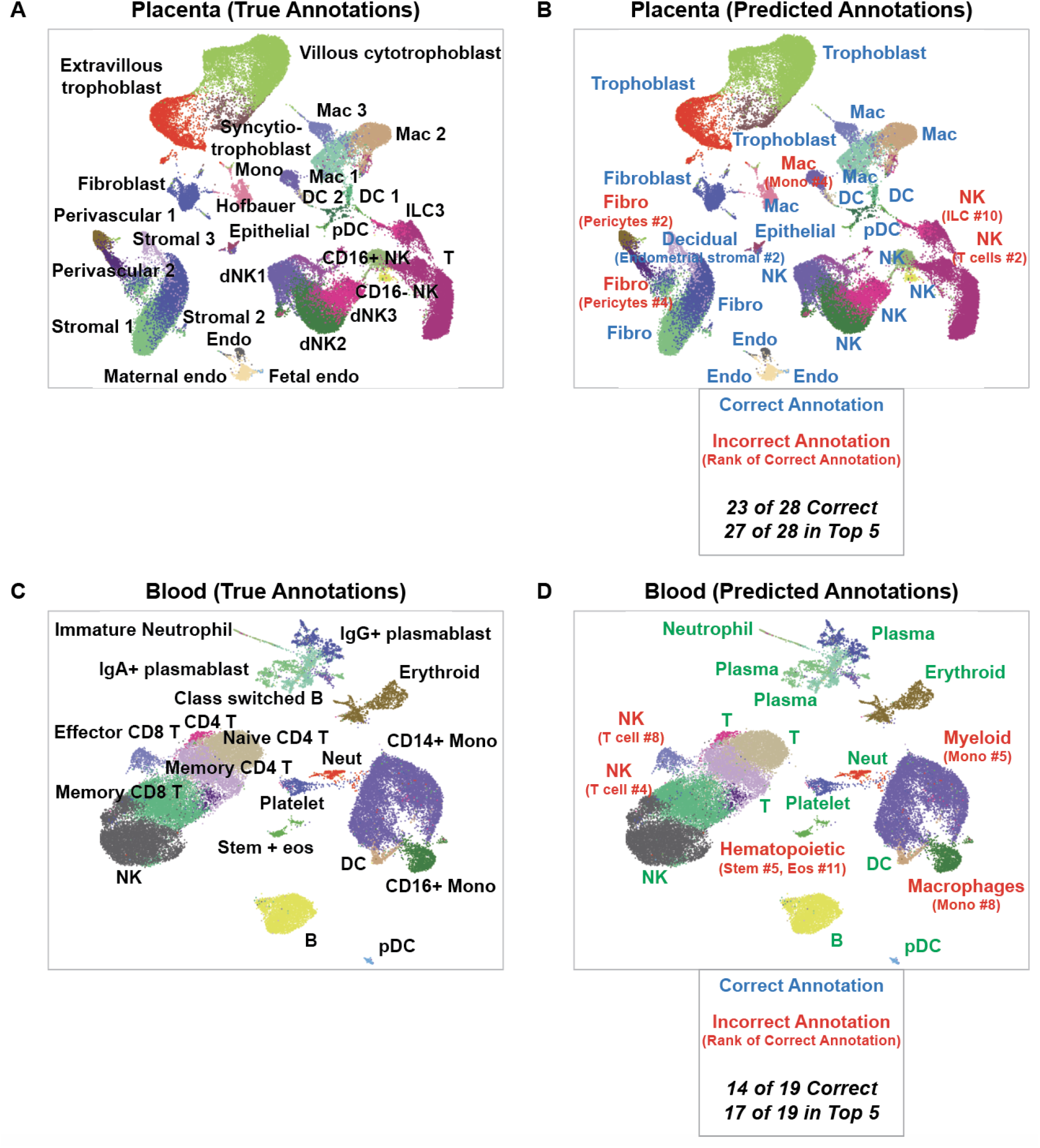
Comparison of true and predicted annotations for all cell types from scRNA-seq studies of the placenta and peripheral blood. True cluster annotations shown in the panels on the left (A, C) are derived from publicly deposited metadata and manual review. Predicted annotations on the right (B, D) are derived from our literature based cell type annotation algorithm, with the optimized parameter settings as described in the text (pan-study reference, top 20 CDGs, scaled local score, weighting by log2FC, and rank by L2 norm). Correct annotations (annotations in which the true priority node exactly matches the mapped priority node for the top prediction) are shown in green, and incorrect annotations are shown in red. For each incorrect annotation, the rank of the correct prediction (out of 103 candidate cell type priority nodes) is shown in parentheses. The datasets selected for display here were previously published in separate studies [28, 29].

**Supplemental Figure 7.**
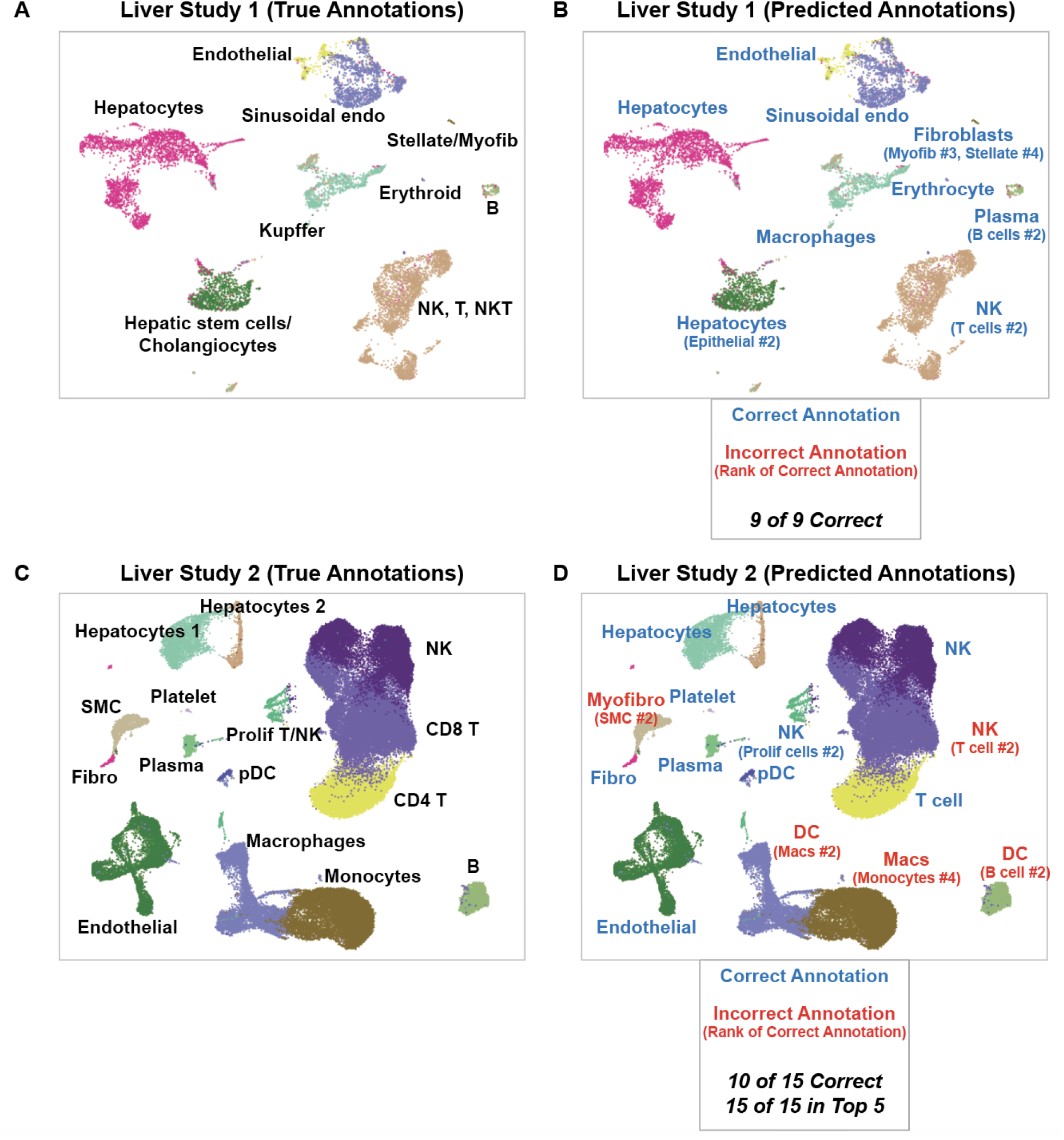
Comparison of true and predicted annotations for all cell types from two scRNA-seq studies of the human liver. True cluster annotations shown in the panels on the left (A, C) are derived from publicly deposited metadata and manual review. Predicted annotations on the right (B, D) are derived from our literature based cell type annotation algorithm, with the optimized parameter settings as described in the text (pan-study reference, top 20 CDGs, scaled local score, weighting by log2FC, and rank by L2 norm). Correct annotations (annotations in which the true priority node exactly matches the mapped priority node for the top prediction) are shown in green, and incorrect annotations are shown in red. For each incorrect annotation, the rank of the correct prediction (out of 103 candidate cell type priority nodes) is shown in parentheses. The datasets selected for display here were previously published in separate studies [27, 37].

**Supplemental Figure 8.**
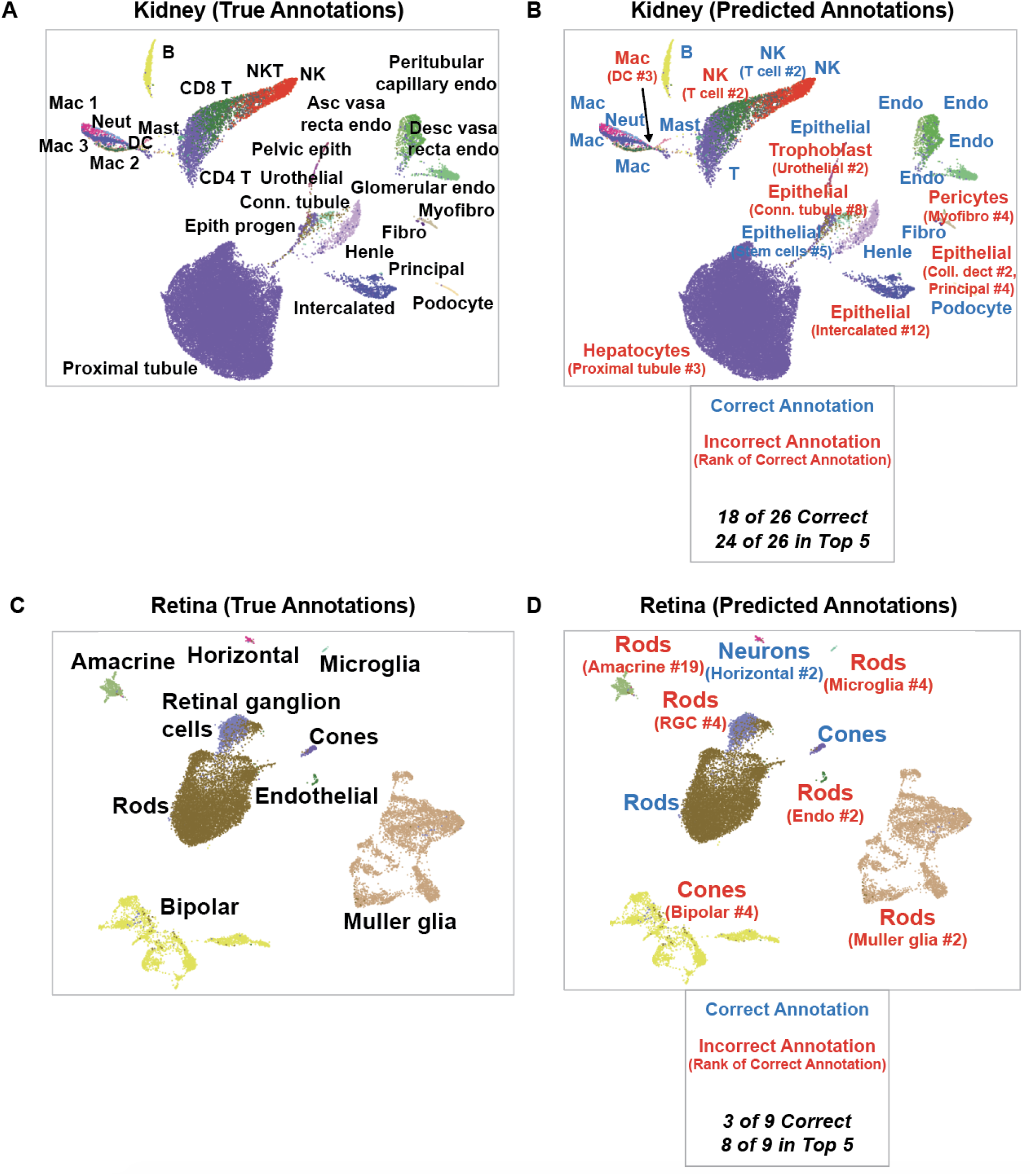
Comparison of true and predicted annotations for all cell types from scRNA-seq studies of the kidney and retina. True cluster annotations shown in the panels on the left (A, C) are derived from publicly deposited metadata and manual review. Predicted annotations on the right (B, D) are derived from our literature based cell type annotation algorithm, with the optimized parameter settings as described in the text (pan-study reference, top 20 CDGs, scaled local score, weighting by log2FC, and rank by L2 norm). Correct annotations (annotations in which the true priority node exactly matches the mapped priority node for the top prediction) are shown in green, and incorrect annotations are shown in red. For each incorrect annotation, the rank of the correct prediction (out of 103 candidate cell type priority nodes) is shown in parentheses. The datasets selected for display here were previously published in separate studies [36, 41].

**Supplemental Figure 9.**
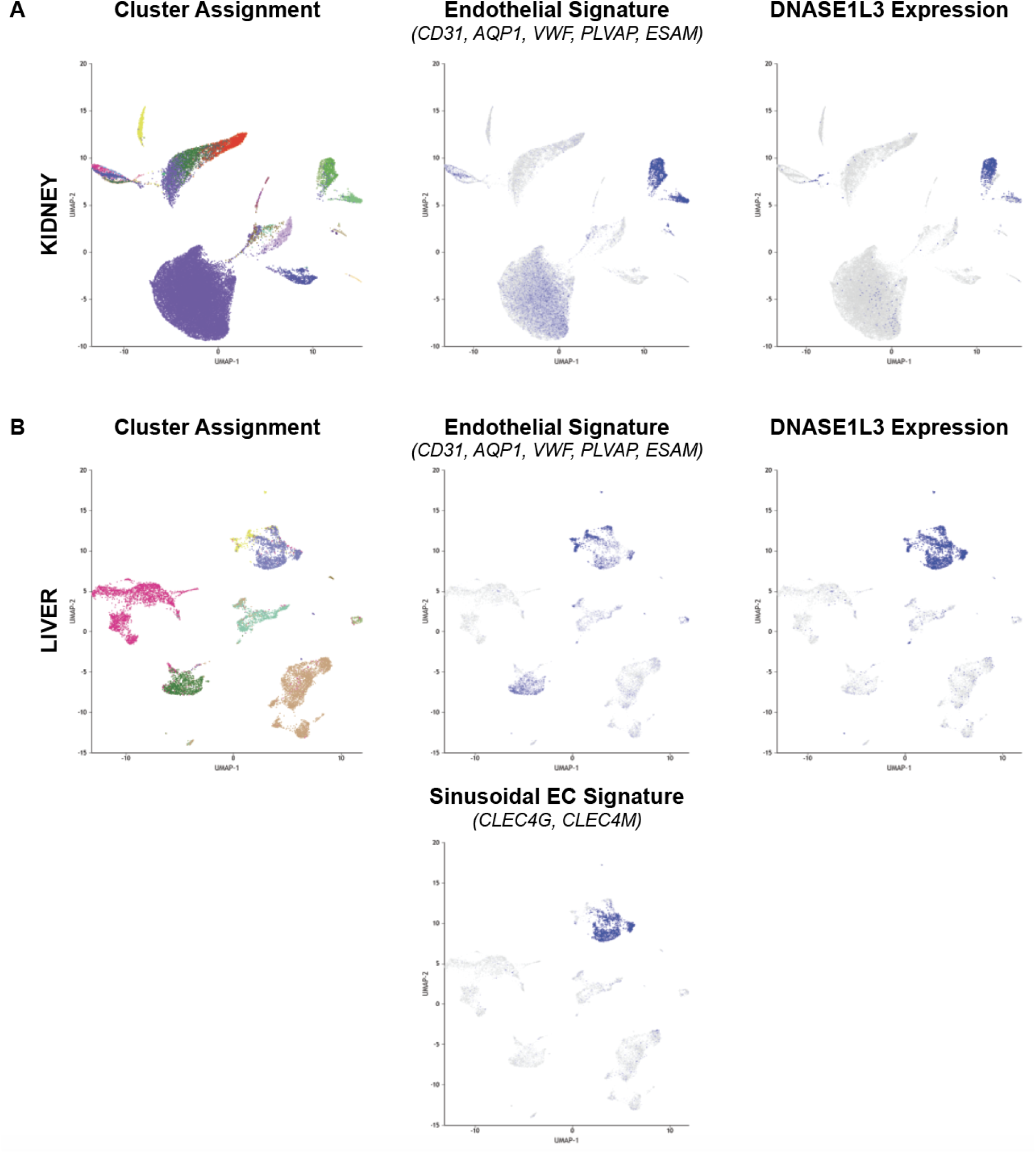
Expression of DNASE1L3 in endothelial cell populations from the kidney and liver. In each panel, the UMAP plot on the far left displays the clusters colored by their cell type annotations; the middle feature plot displays the expression level of a gene signature comprised of five canonical endothelial markers (CD31, AQP1, VWF, PLVAP, and ESAM) or two sinusoidal endothelial cell markers (CLEC4G and CLEC4M); and the far right feature plot displays the expression level of DNASE1L3, which overlaps with the endothelial signatures in each case. The data is derived from (A) kidney [36] and (B) liver [27].

**Supplemental Figure 10.**
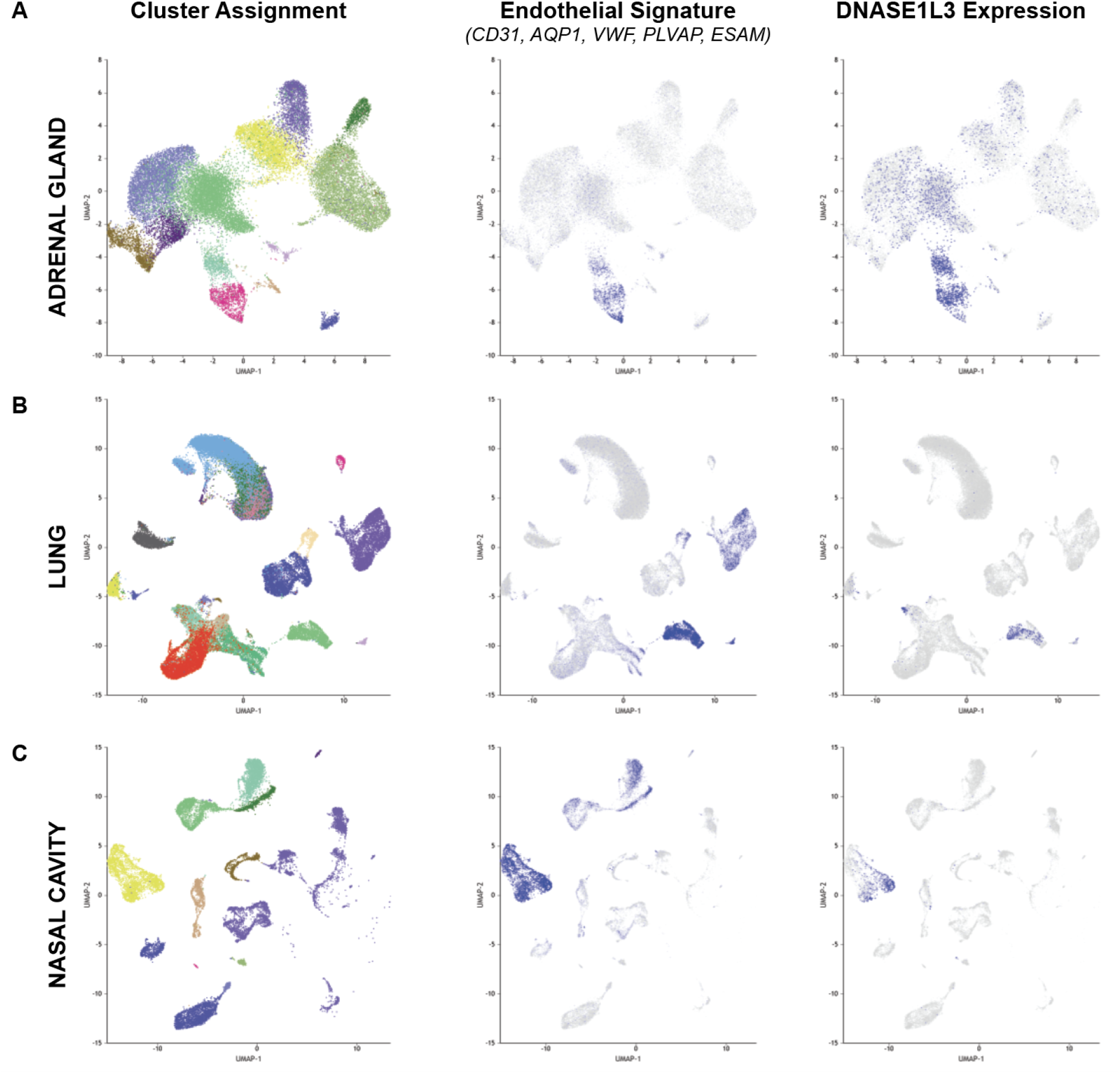
Expression of DNASE1L3 in endothelial cells from adrenal gland, lung, and nasal cavity. In each panel, the UMAP plot on the far left displays the clusters colored by their cell type annotations; the middle feature plot displays the expression level of a gene signature comprised of five canonical endothelial markers (CD31, AQP1, VWF, PLVAP, and ESAM); and the far right feature plot displays the expression level of DNASE1L3, which overlaps with the endothelial signature in each case. The data is derived from (A) adrenal gland [53], (B) lung [44], and (c) nasal cavity [51].

## Supplemental Tables

**Supplemental Table 1.**
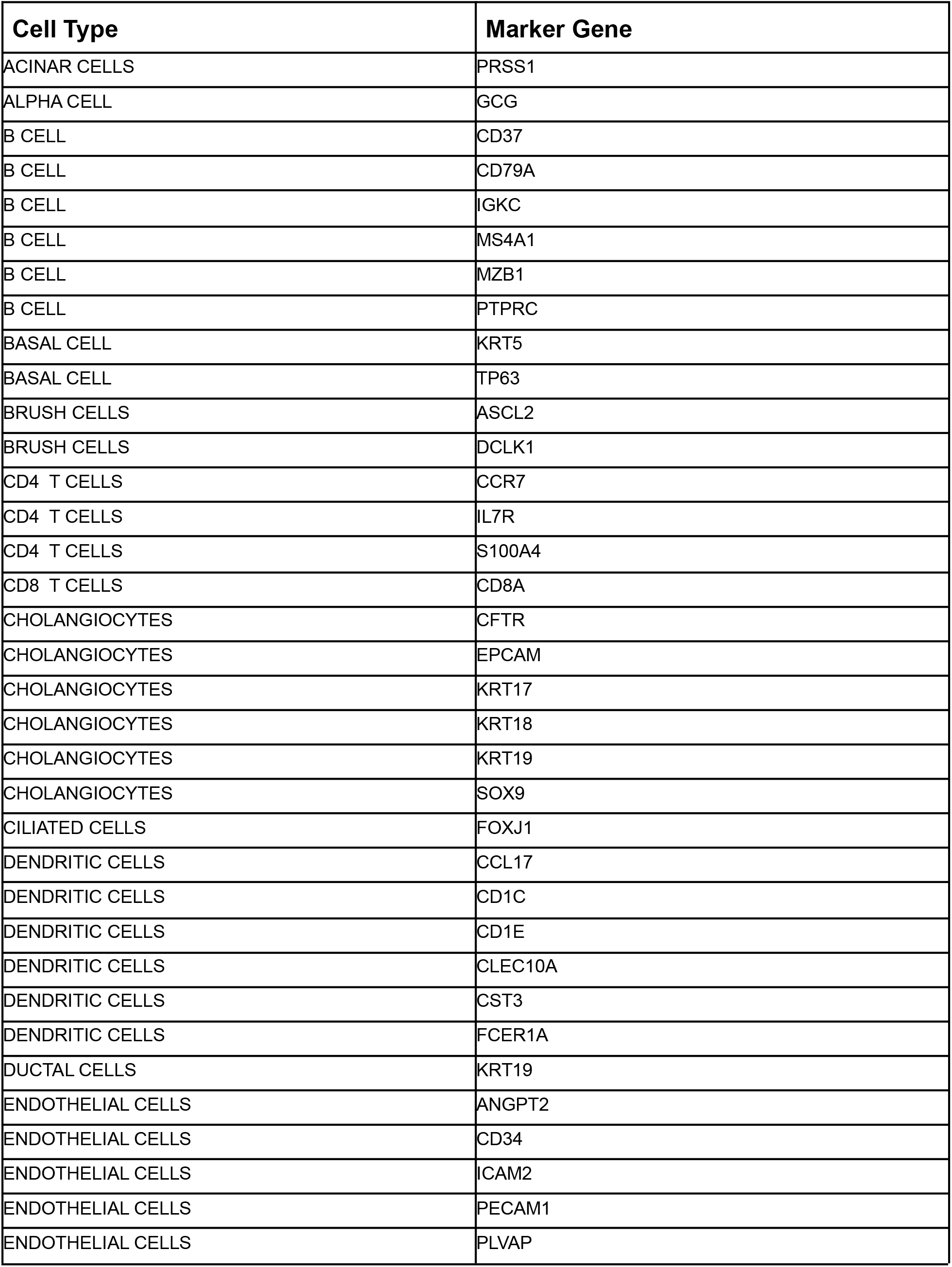
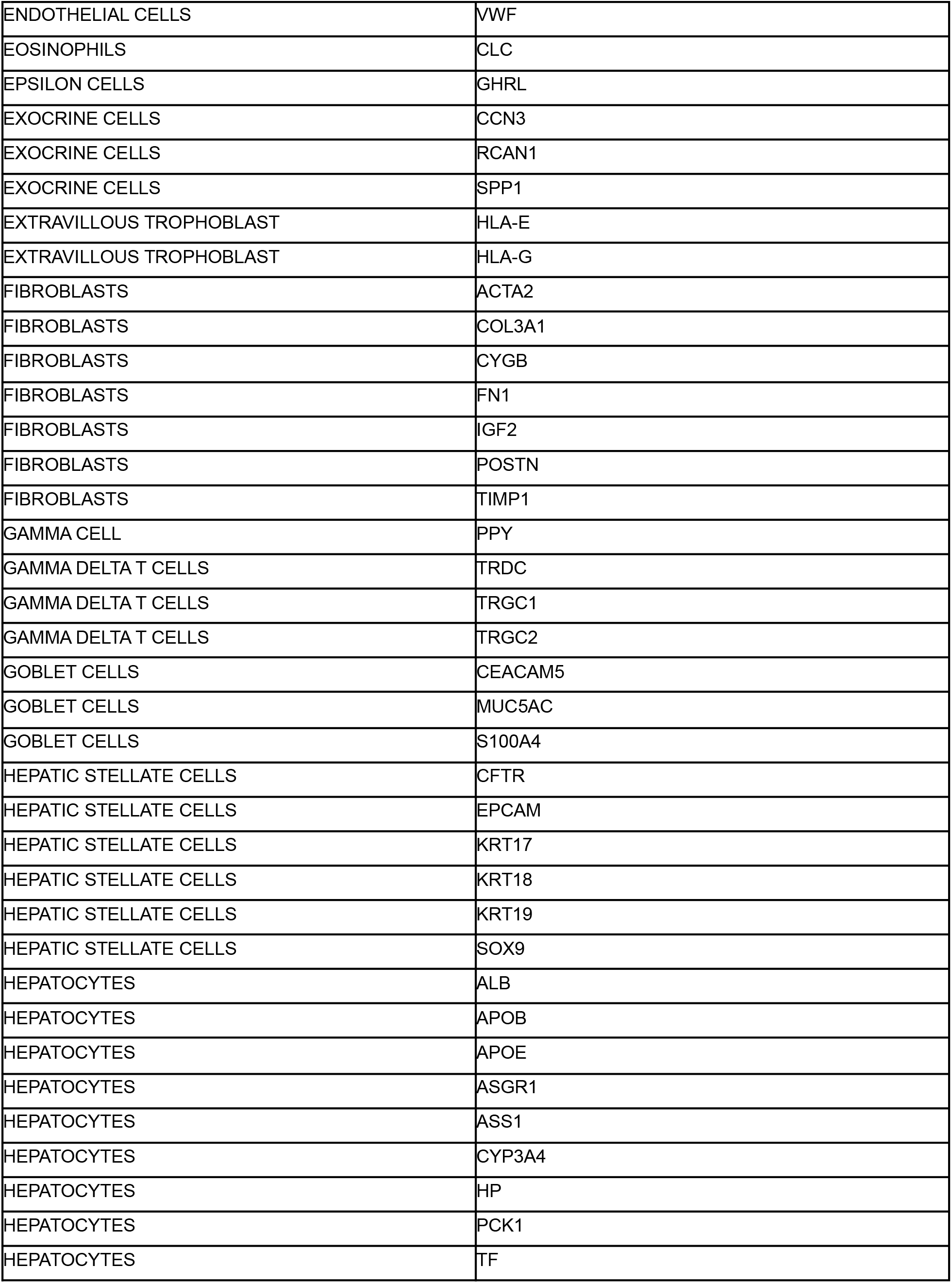
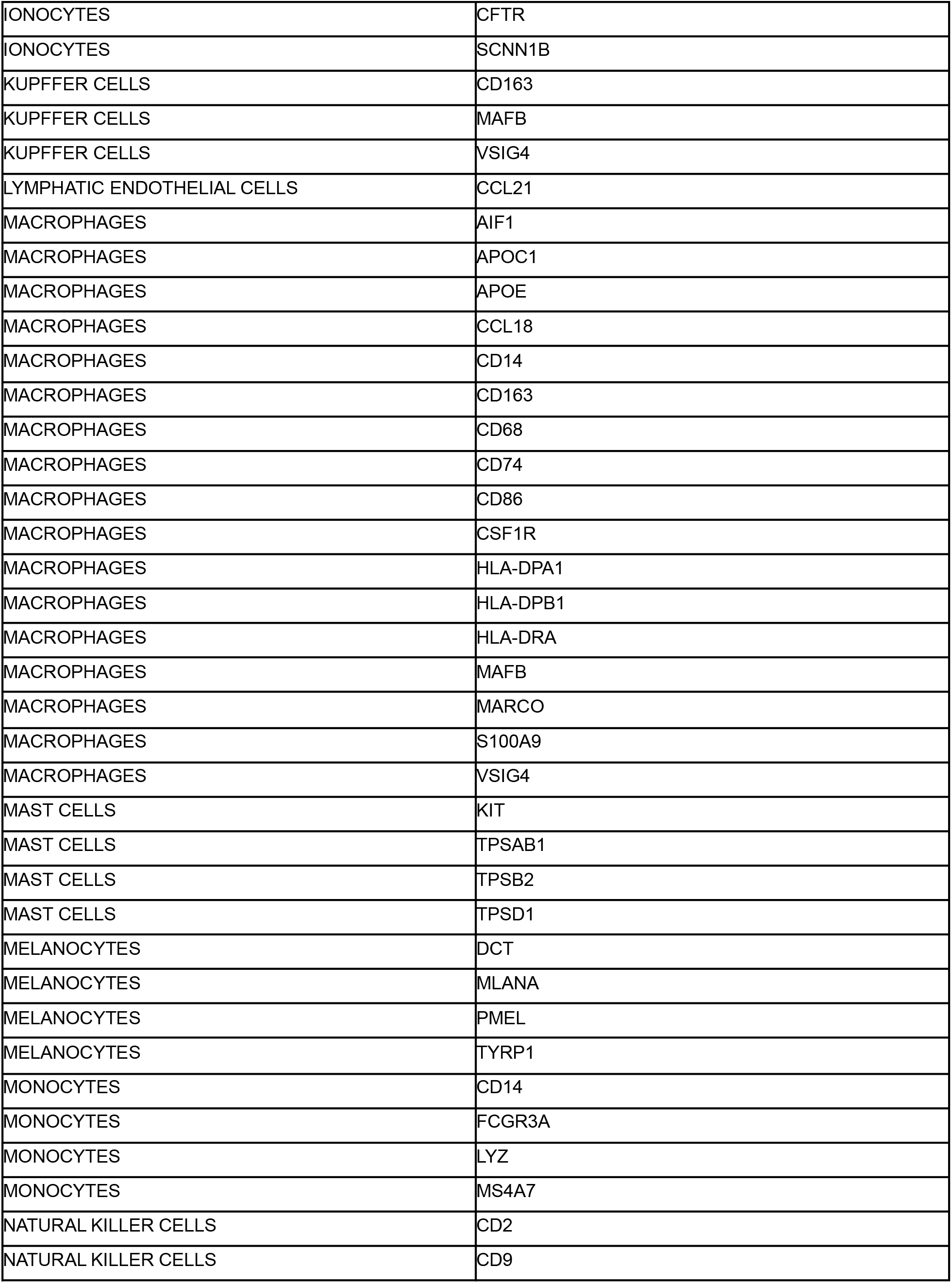
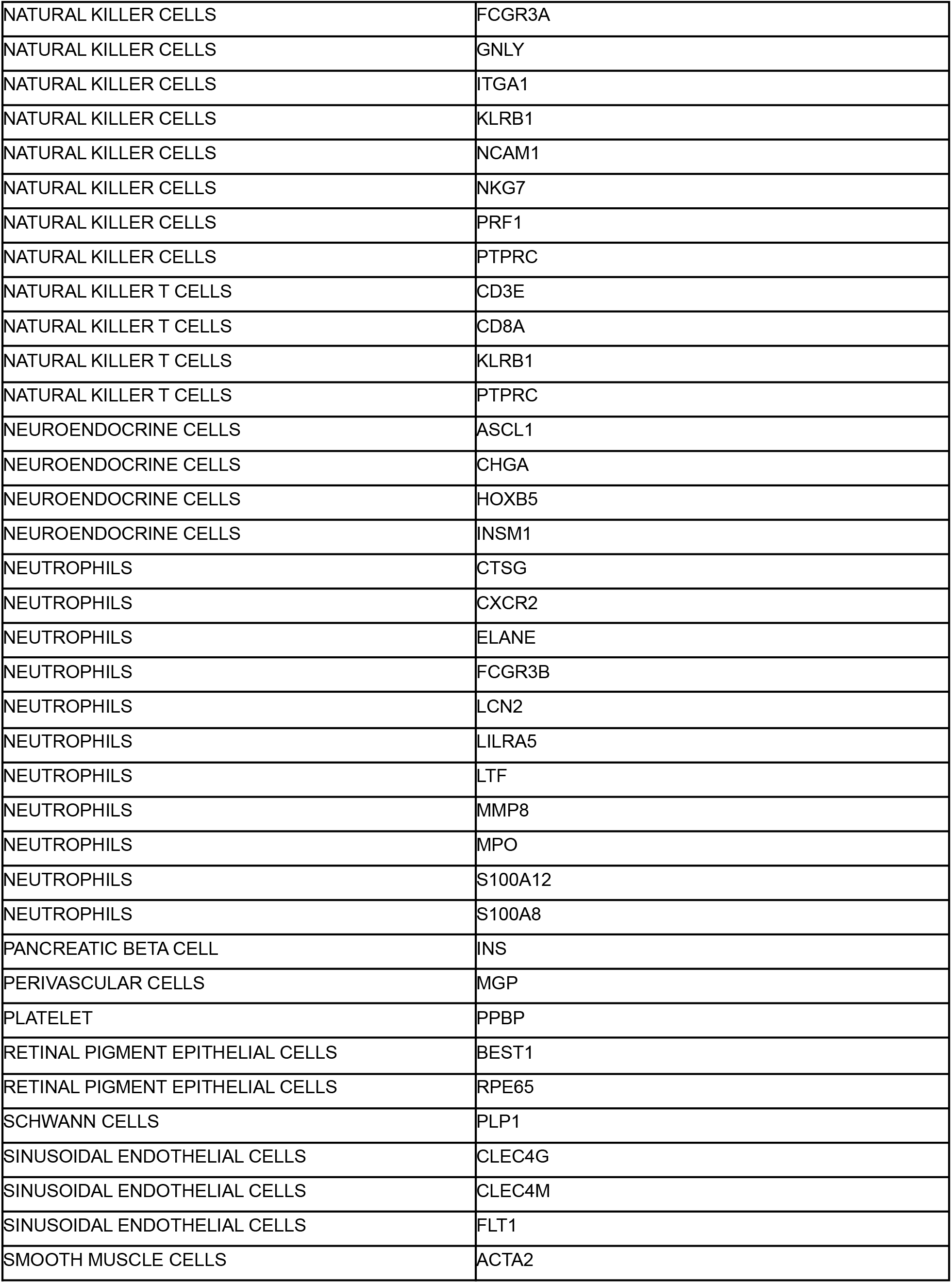
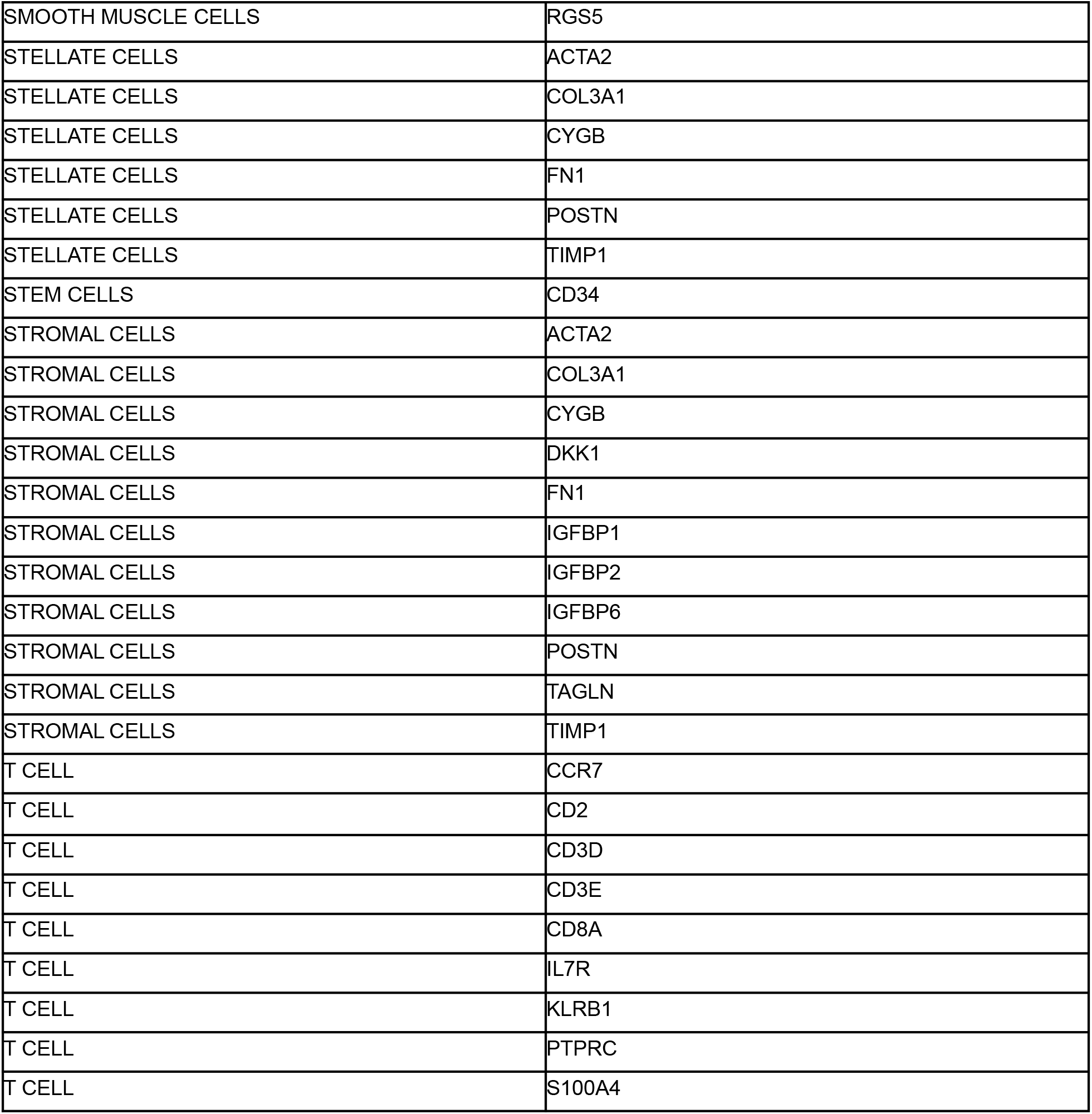
Manually curated cell type defining genes. 174 gene-cell type pairs were extracted from published scRNA-seq datasets in which marker genes that were used for manual cluster annotation were reported [25–32]. This set of gene-cell type pairs were designated as “matched”, and all other possible pairwise combinations of these genes and cell types were designated as “mismatched” pairs for subsequent analyses.

| **Cell Type** | **Marker Gene** |
| --- | --- |
| ACINAR CELLS | PRSS1 |
| ALPHA CELL | GCG |
| B CELL | CD37 |
| B CELL | CD79A |
| B CELL | IGKC |
| B CELL | MS4A1 |
| B CELL | MZB1 |
| B CELL | PTPRC |
| BASAL CELL | KRT5 |
| BASAL CELL | TP63 |
| BRUSH CELLS | ASCL2 |
| BRUSH CELLS | DCLK1 |
| CD4 T CELLS | CCR7 |
| CD4 T CELLS | IL7R |
| CD4 T CELLS | S100A4 |
| CD8 T CELLS | CD8A |
| CHOLANGIOCYTES | CFTR |
| CHOLANGIOCYTES | EPCAM |
| CHOLANGIOCYTES | KRT17 |
| CHOLANGIOCYTES | KRT18 |
| CHOLANGIOCYTES | KRT19 |
| CHOLANGIOCYTES | SOX9 |
| CILIATED CELLS | FOXJ1 |
| DENDRITIC CELLS | CCL17 |
| DENDRITIC CELLS | CD1C |
| DENDRITIC CELLS | CD1E |
| DENDRITIC CELLS | CLEC10A |
| DENDRITIC CELLS | CST3 |
| DENDRITIC CELLS | FCER1A |
| DUCTAL CELLS | KRT19 |
| ENDOTHELIAL CELLS | ANGPT2 |
| ENDOTHELIAL CELLS | CD34 |
| ENDOTHELIAL CELLS | ICAM2 |
| ENDOTHELIAL CELLS | PECAM1 |
| ENDOTHELIAL CELLS | PLVAP |
| ENDOTHELIAL CELLS | VWF |
| EOSINOPHILS | CLC |
| EPSILON CELLS | GHRL |
| EXOCRINE CELLS | CCN3 |
| EXOCRINE CELLS | RCAN1 |
| EXOCRINE CELLS | SPP1 |
| EXTRAVILLOUS TROPHOBLAST | HLA-E |
| EXTRAVILLOUS TROPHOBLAST | HLA-G |
| FIBROBLASTS | ACTA2 |
| FIBROBLASTS | COL3A1 |
| FIBROBLASTS | CYGB |
| FIBROBLASTS | FN1 |
| FIBROBLASTS | IGF2 |
| FIBROBLASTS | POSTN |
| FIBROBLASTS | TIMP1 |
| GAMMA CELL | PPY |
| GAMMA DELTA T CELLS | TRDC |
| GAMMA DELTA T CELLS | TRGC1 |
| GAMMA DELTA T CELLS | TRGC2 |
| GOBLET CELLS | CEACAM5 |
| GOBLET CELLS | MUC5AC |
| GOBLET CELLS | S100A4 |
| HEPATIC STELLATE CELLS | CFTR |
| HEPATIC STELLATE CELLS | EPCAM |
| HEPATIC STELLATE CELLS | KRT17 |
| HEPATIC STELLATE CELLS | KRT18 |
| HEPATIC STELLATE CELLS | KRT19 |
| HEPATIC STELLATE CELLS | SOX9 |
| HEPATOCYTES | ALB |
| HEPATOCYTES | APOB |
| HEPATOCYTES | APOE |
| HEPATOCYTES | ASGR1 |
| HEPATOCYTES | ASS1 |
| HEPATOCYTES | CYP3A4 |
| HEPATOCYTES | HP |
| HEPATOCYTES | PCK1 |
| HEPATOCYTES | TF |
| IONOCYTES | CFTR |
| IONOCYTES | SCNN1B |
| KUPFFER CELLS | CD163 |
| KUPFFER CELLS | MAFB |
| KUPFFER CELLS | VSIG4 |
| LYMPHATIC ENDOTHELIAL CELLS | CCL21 |
| MACROPHAGES | AIF1 |
| MACROPHAGES | APOC1 |
| MACROPHAGES | APOE |
| MACROPHAGES | CCL18 |
| MACROPHAGES | CD14 |
| MACROPHAGES | CD163 |
| MACROPHAGES | CD68 |
| MACROPHAGES | CD74 |
| MACROPHAGES | CD86 |
| MACROPHAGES | CSF1R |
| MACROPHAGES | HLA-DPA1 |
| MACROPHAGES | HLA-DPB1 |
| MACROPHAGES | HLA-DRA |
| MACROPHAGES | MAFB |
| MACROPHAGES | MARCO |
| MACROPHAGES | S100A9 |
| MACROPHAGES | VSIG4 |
| MAST CELLS | KIT |
| MAST CELLS | TPSAB1 |
| MAST CELLS | TPSB2 |
| MAST CELLS | TPSD1 |
| MELANOCYTES | DCT |
| MELANOCYTES | MLANA |
| MELANOCYTES | PMEL |
| MELANOCYTES | TYRP1 |
| MONOCYTES | CD14 |
| MONOCYTES | FCGR3A |
| MONOCYTES | LYZ |
| MONOCYTES | MS4A7 |
| NATURAL KILLER CELLS | CD2 |
| NATURAL KILLER CELLS | CD9 |
| NATURAL KILLER CELLS | FCGR3A |
| NATURAL KILLER CELLS | GNLY |
| NATURAL KILLER CELLS | ITGA1 |
| NATURAL KILLER CELLS | KLRB1 |
| NATURAL KILLER CELLS | NCAM1 |
| NATURAL KILLER CELLS | NKG7 |
| NATURAL KILLER CELLS | PRF1 |
| NATURAL KILLER CELLS | PTPRC |
| NATURAL KILLER T CELLS | CD3E |
| NATURAL KILLER T CELLS | CD8A |
| NATURAL KILLER T CELLS | KLRB1 |
| NATURAL KILLER T CELLS | PTPRC |
| NEUROENDOCRINE CELLS | ASCL1 |
| NEUROENDOCRINE CELLS | CHGA |
| NEUROENDOCRINE CELLS | HOXB5 |
| NEUROENDOCRINE CELLS | INSM1 |
| NEUTROPHILS | CTSG |
| NEUTROPHILS | CXCR2 |
| NEUTROPHILS | ELANE |
| NEUTROPHILS | FCGR3B |
| NEUTROPHILS | LCN2 |
| NEUTROPHILS | LILRA5 |
| NEUTROPHILS | LTF |
| NEUTROPHILS | MMP8 |
| NEUTROPHILS | MPO |
| NEUTROPHILS | S100A12 |
| NEUTROPHILS | S100A8 |
| PANCREATIC BETA CELL | INS |
| PERIVASCULAR CELLS | MGP |
| PLATELET | PPBP |
| RETINAL PIGMENT EPITHELIAL CELLS | BEST1 |
| RETINAL PIGMENT EPITHELIAL CELLS | RPE65 |
| SCHWANN CELLS | PLP1 |
| SINUSOIDAL ENDOTHELIAL CELLS | CLEC4G |
| SINUSOIDAL ENDOTHELIAL CELLS | CLEC4M |
| SINUSOIDAL ENDOTHELIAL CELLS | FLT1 |
| SMOOTH MUSCLE CELLS | ACTA2 |
| SMOOTH MUSCLE CELLS | RGS5 |
| STELLATE CELLS | ACTA2 |
| STELLATE CELLS | COL3A1 |
| STELLATE CELLS | CYGB |
| STELLATE CELLS | FN1 |
| STELLATE CELLS | POSTN |
| STELLATE CELLS | TIMP1 |
| STEM CELLS | CD34 |
| STROMAL CELLS | ACTA2 |
| STROMAL CELLS | COL3A1 |
| STROMAL CELLS | CYGB |
| STROMAL CELLS | DKK1 |
| STROMAL CELLS | FN1 |
| STROMAL CELLS | IGFBP1 |
| STROMAL CELLS | IGFBP2 |
| STROMAL CELLS | IGFBP6 |
| STROMAL CELLS | POSTN |
| STROMAL CELLS | TAGLN |
| STROMAL CELLS | TIMP1 |
| T CELL | CCR7 |
| T CELL | CD2 |
| T CELL | CD3D |
| T CELL | CD3E |
| T CELL | CD8A |
| T CELL | IL7R |
| T CELL | KLRB1 |
| T CELL | PTPRC |
| T CELL | S100A4 |

**Supplemental Table 2.**
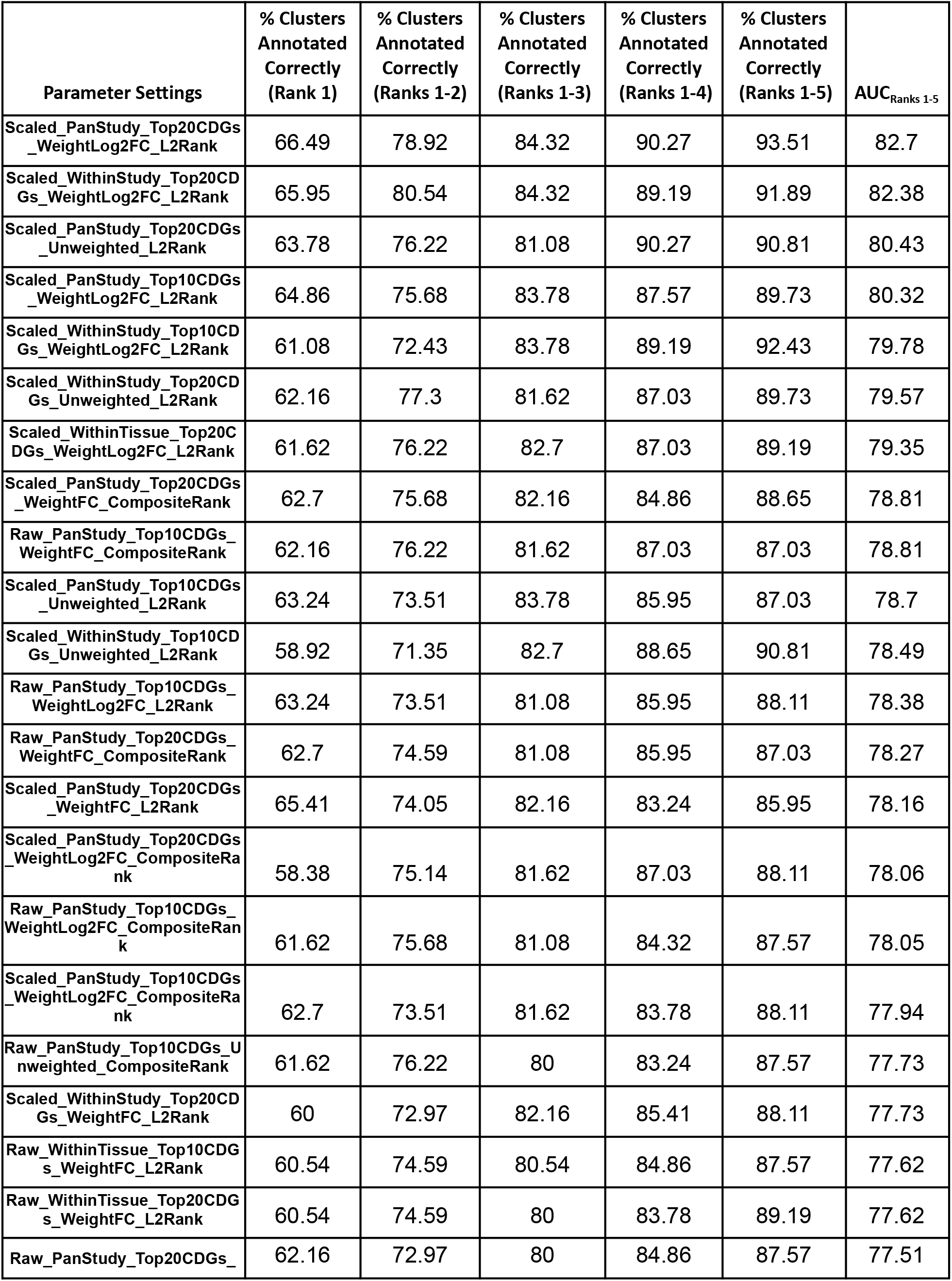
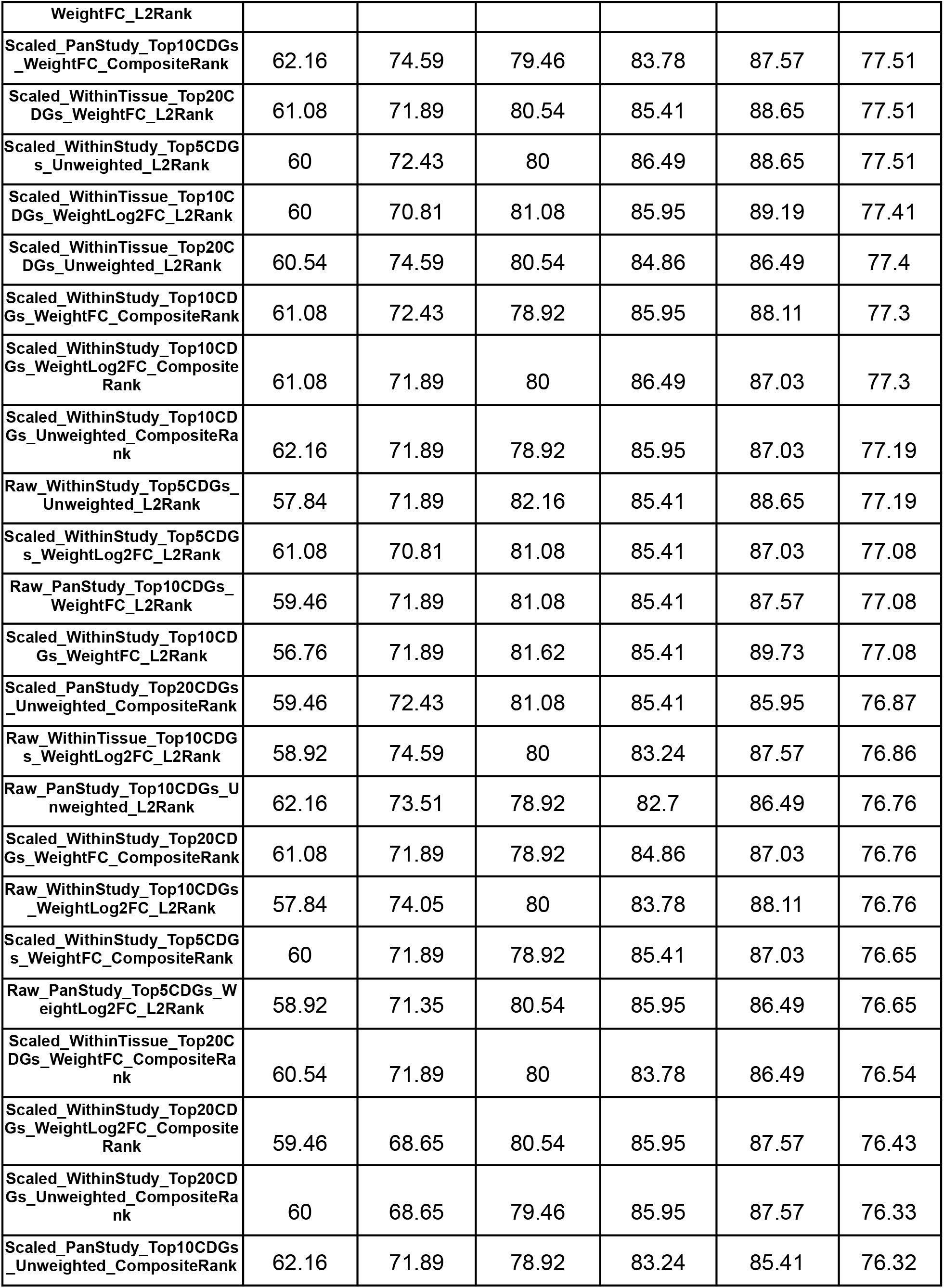
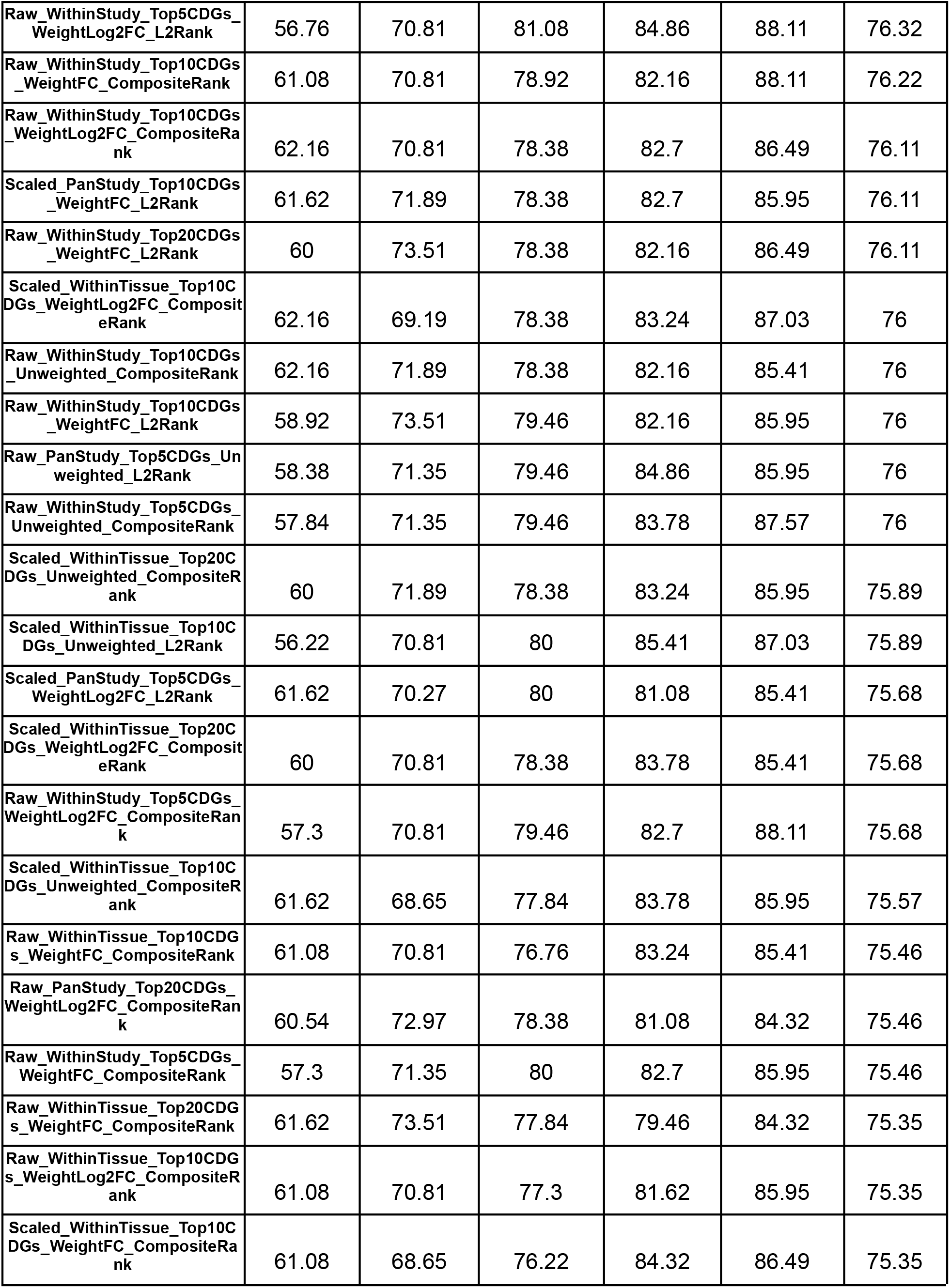
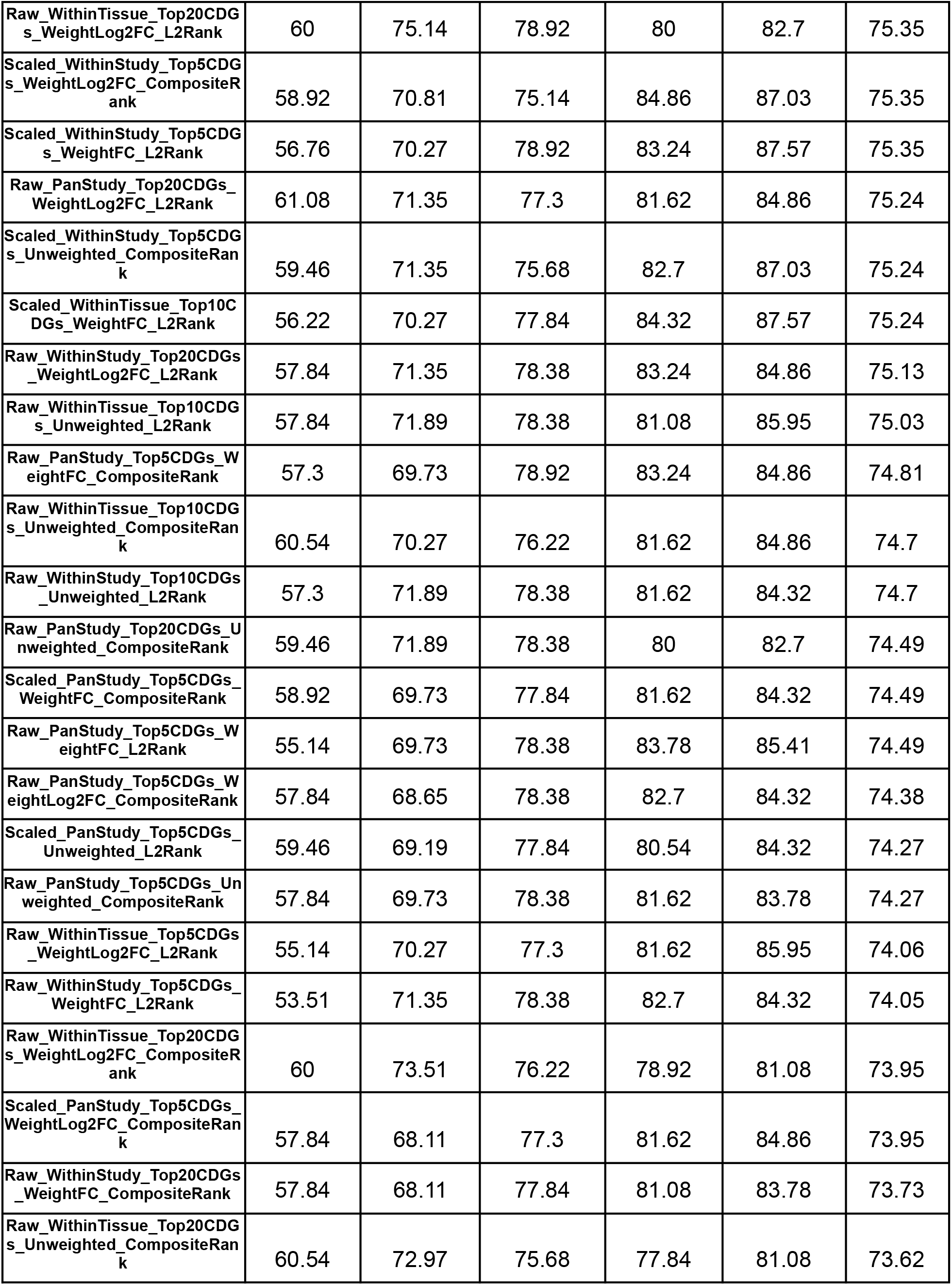
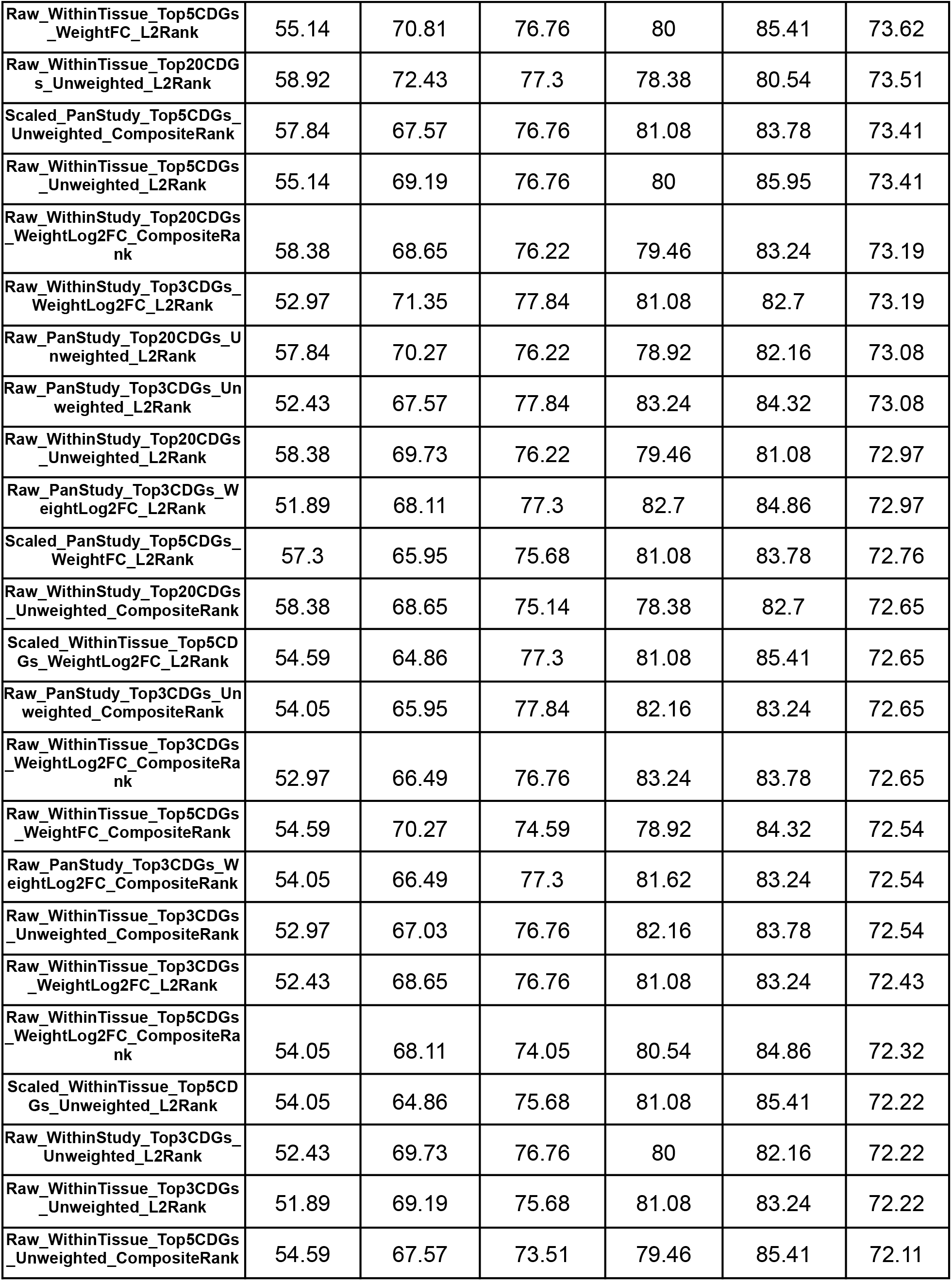
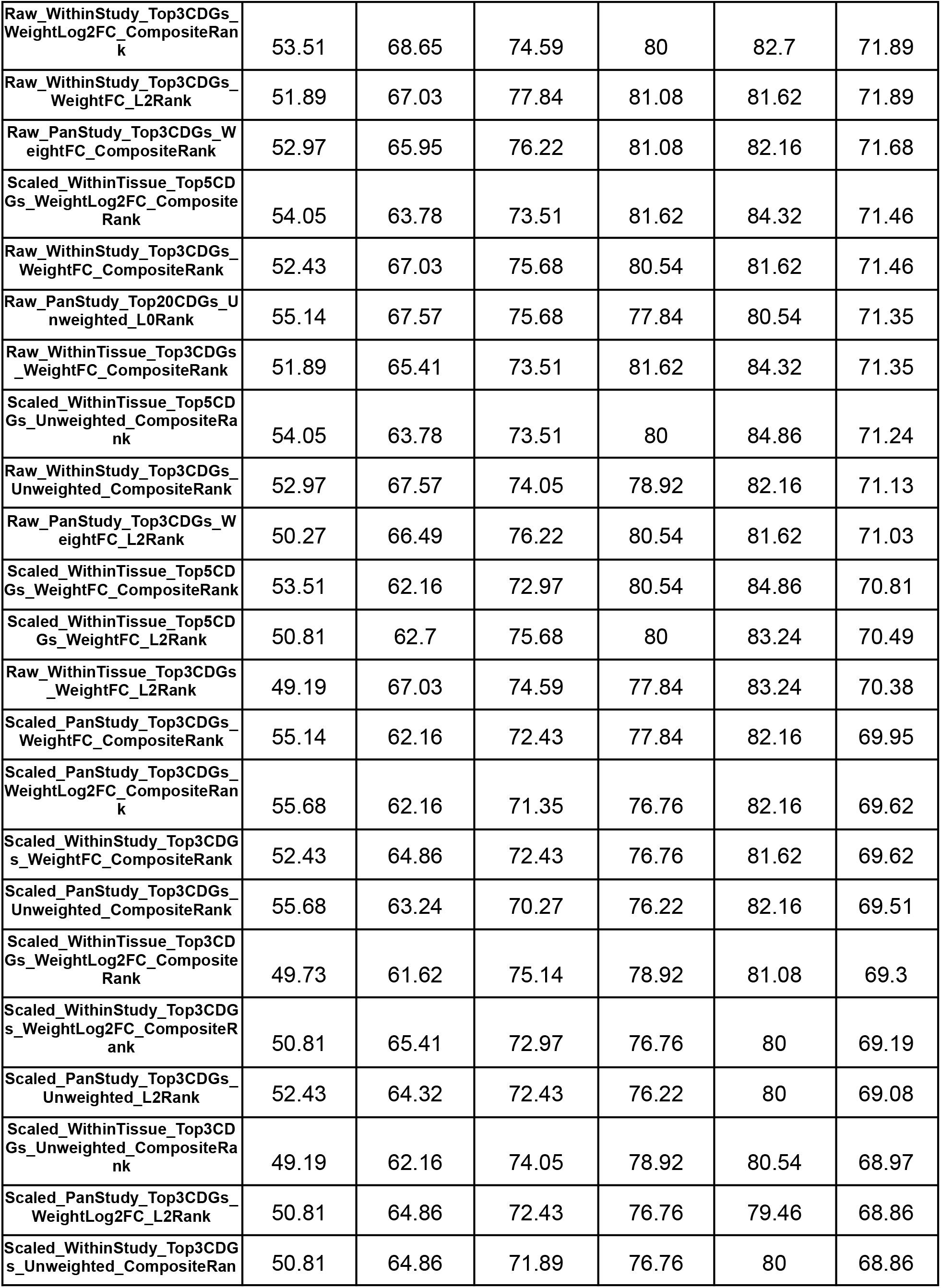
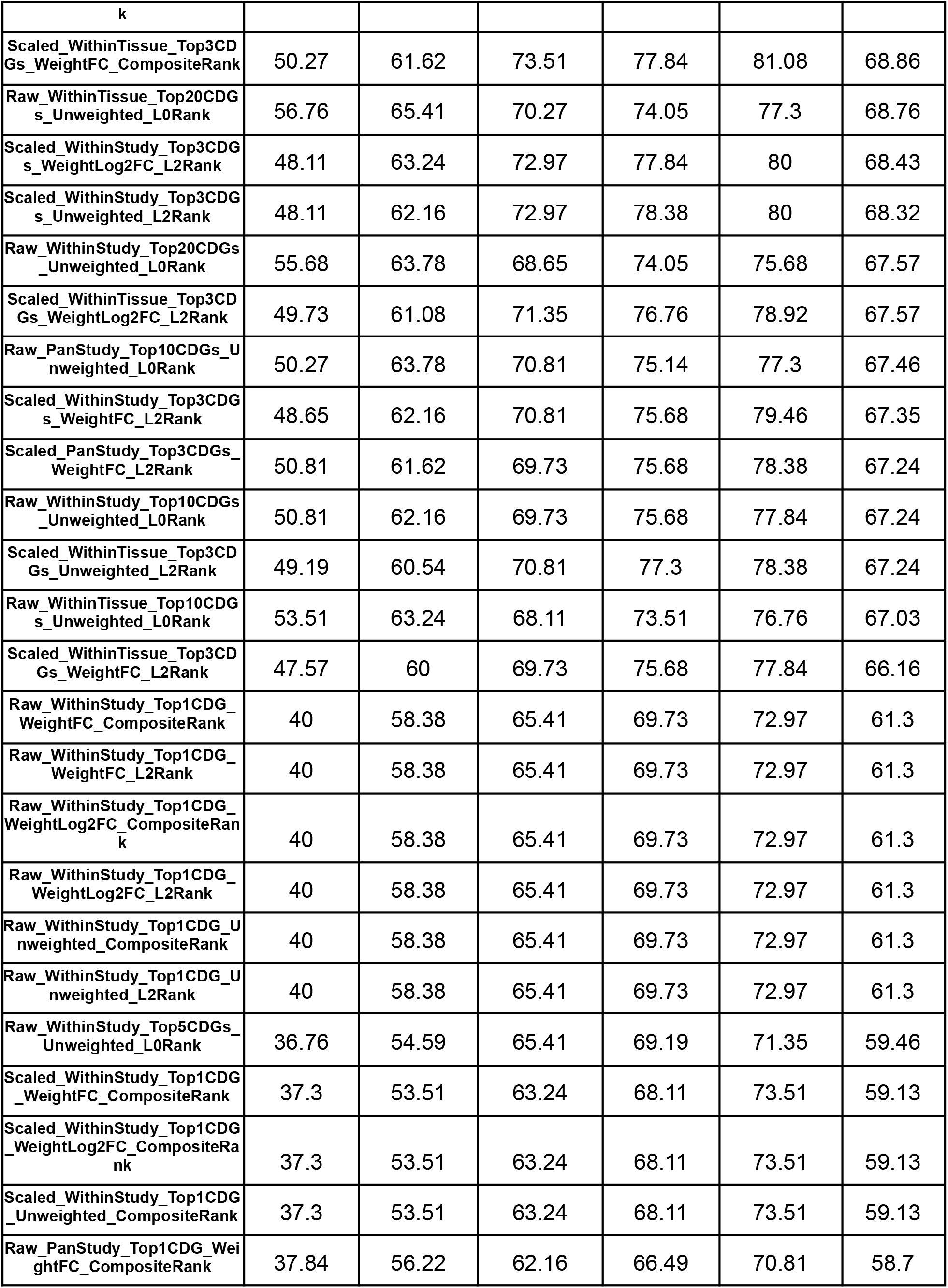
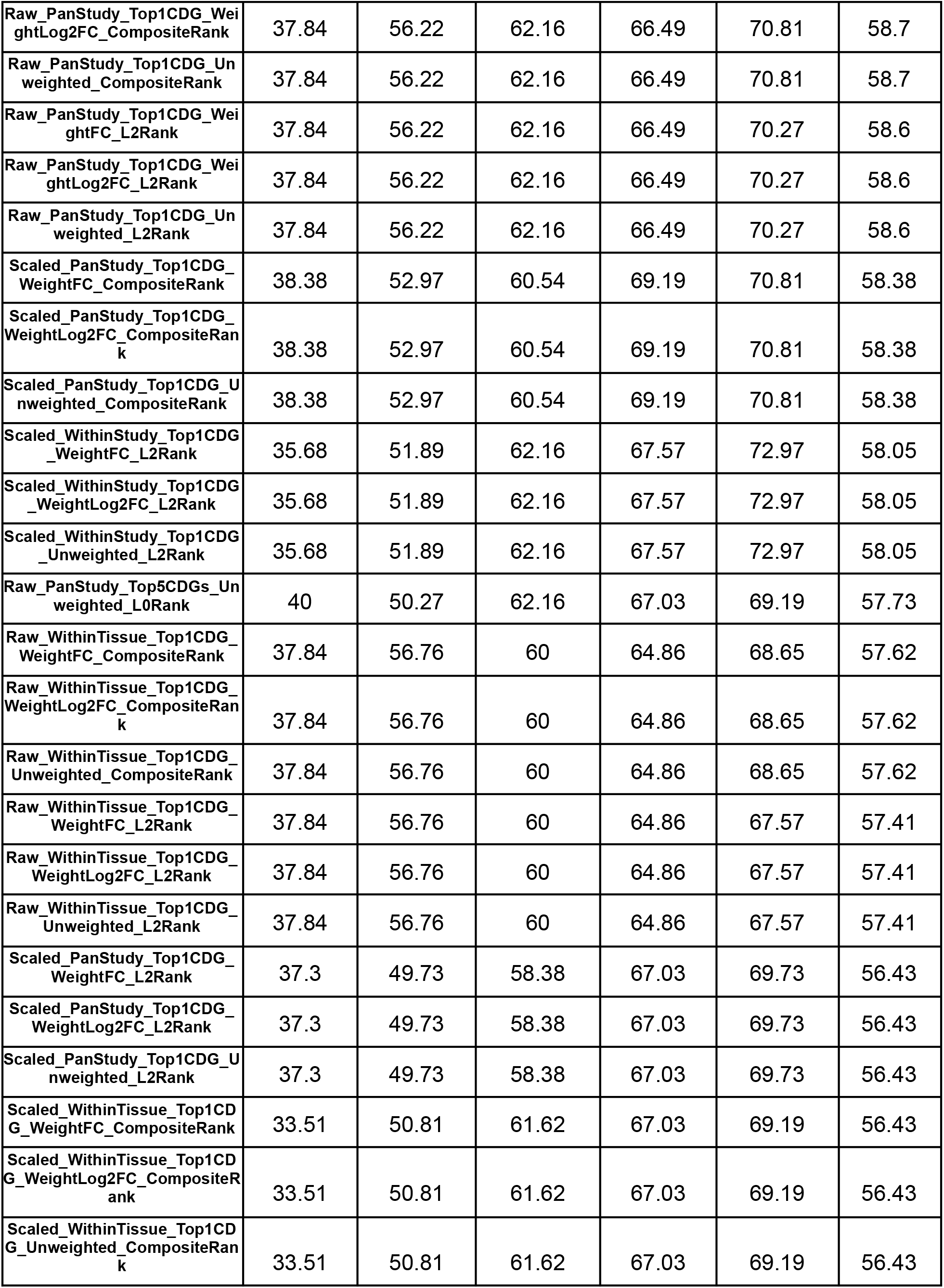
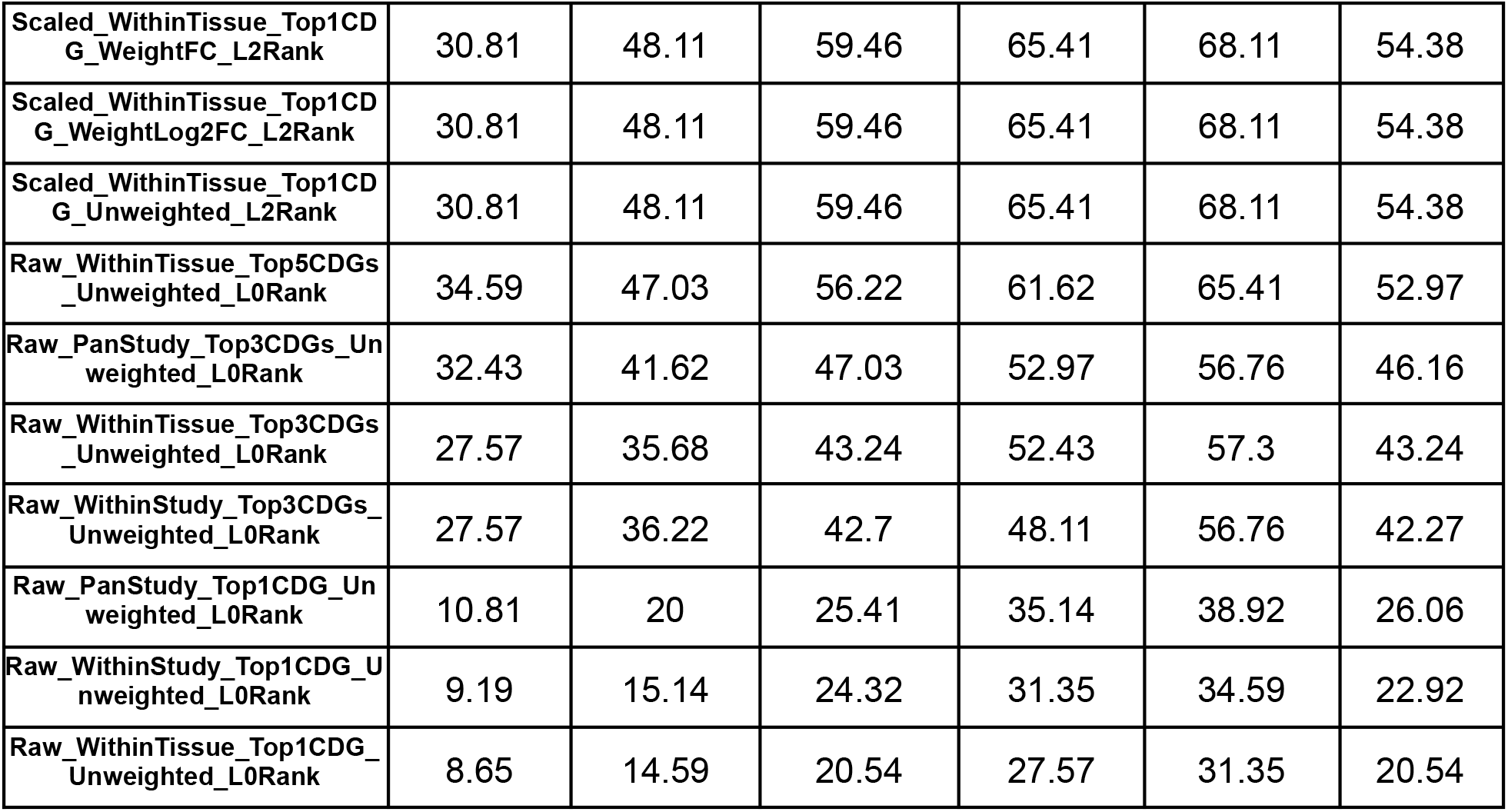
Summary of annotation algorithm performance for all tested parameter combinations. The “Parameter Settings” column indicates the combination of parameters which was used to annotate clusters, including the following listed in this order: (1) local score version (raw or scaled), (2) reference cells used to calculate CDGs (pan-study, within tissue, or within study), (3) number of CDGs used (1, 3, 5, 10, or 20), (4) the weighting method used calculating L2 norms (unweighted, fold change, or log2FC), and (5) the ranking metric used (modified L0 rank, L2 rank, or composite rank). The next five columns indicate the percentage of clusters (out of 185) for which the correct annotation was among the top-ranked 1, 2, 3, 4, or 5 predictions. The last column provides the AUC_Ranks 1-5_ metric for each parameter combination, which was calculated as the average of the previous five columns. The table is sorted in descending order by AUC_Ranks 1-5_, which was used to assess overall algorithm performance.

